# Astrocyte senescence impairs synaptogenesis due to Thrombospondin-1 loss

**DOI:** 10.64898/2025.12.02.691780

**Authors:** Stefano Ercoli, Lucía Casares-Crespo, Elena Juárez-Escoto, Helena Mira

## Abstract

Cellular senescence is an irreversible state linked to aging that involves molecular and functional alterations. The mammalian hippocampus, a key brain region for learning and memory, is highly vulnerable to damage in age-related neurodegenerative diseases, yet the role of cellular senescence in hippocampal aging remains underexplored. Here, we report an early onset of senescence signatures in hippocampal astrocytes of the accelerated aging and frailty mouse model SAMP8. We examine how astrocyte senescence affects excitatory synapse formation, focusing on soluble signals released by astrocytes. Astrocytes isolated from SAMP8 brain and those differentiated from SAMP8 neural stem cells show senescence hallmarks (SA-β-gal, p16^INK4a^, Lamin B1 loss), alongside a significant reduction in synaptogenic function. While astrocyte-conditioned medium (ACM) from control mice promotes excitatory synaptogenesis through thrombospondin-1 / α2δ-1 neuronal receptor signalling, ACM from senescent SAMP8 astrocytes lacks this capacity. Supplementing senescent ACM with thrombospondin-1 protein, or overexpressing thrombospondin-1 gene in senescent astrocytes, reinstates synaptogenesis. At the hippocampal level, thrombospondin-1 and synaptic puncta are reduced in SAMP8 mice. Our findings reveal that senescent astrocytes exhibit reduced synaptogenic capacity due to thrombospondin-1 loss, highlighting their contribution to synaptic dysfunction during aging. Preventing senescence in hippocampal astrocytes may thus restore astrocyte-mediated synaptogenesis in the aged brain.

## Introduction

Aging is a complex process that involves the progressive deterioration of the organism. It is characterized by the accumulation of damage throughout life and the gradual loss of tissue and organ function (Kirkwood, 2017). Scientific discoveries in recent decades have identified a set of interconnected processes, known as the “hallmarks of aging,” considered as a starting point for exploring the foundations of aging and for developing treatments to counteract age-related functional decline (López-Otín et al., 2023).

Cellular senescence, a hallmark of aging, stands out among the causal processes of organismal aging (Campisi, 2005). In proliferating cells, senescence has been defined as a state of irreversible cell cycle arrest that involves genetic, morphological, and functional alterations with a marked negative impact on tissue function (Cohen and Torres, 2019; Dos Santos et al., 2024). Both proliferating and non-proliferating cells can undergo senescence in response to a variety of internal and external factors, including cytotoxic stimuli, oxidative stress, proteasome inhibition, DNA damage, and alterations in cellular communication systems, due to paracrine signals released by neighbouring senescent cells (Takai, Smogorzewska, & de Lange, 2003; Torres, Lewis, and Cristofalo, 2006; Pricola et al., 2009; Bitto et al., 2010; Rodier et al., 2011; Acosta et al., 2013; Clevers & Watts, 2018). Cellular senescence can be detected through a variety of senescence markers such as increased expression of the cyclin-dependent kinase (CDK) inhibitors CDKN2A/p16^INK4a^ and CDKN1A/p21^CIP1^ (Evans et al. 2003; Bhat et al., 2012; Yu et al., 2017; Idda et al., 2020), changes in nuclear structure and decreased levels of the nuclear lamina protein Lamin B1 (Freund et al., 2012; Matias et al., 2021), increased lysosomal mass linked to the detection of elevated levels of senescence-associated β-galactosidase activity (SA-β-gal) at pH 6 (Dimri et al., 1995; Evans et al., 2003; Piechota et al., 2016; Yu et al., 2017), and increased secretion of proinflammatory chemokines, cytokines, proteases and growth factors, collectively known as the senescence-associated secretory phenotype (SASP) (Coppé et al., 2008; Bhat et al., 2012; Cohen et al., 2017; Gorgoulis et al., 2019). Despite the close association between cellular senescence and age-related tissue dysfunction, the role of cellular senescence in brain aging remains poorly explored.

The SAMP8 (Senescence-Accelerated Mouse Prone 8) mouse is a non-transgenic model of accelerated aging widely used to study brain aging. Compared to the control SAMR1 (Senescence-Accelerated Mouse Resistant 1) strain, SAMP8 animals show early signs of motor impairment, muscle weakness, and memory deficits, as observed in human aging (Dacomo et al., 2024). At the brain level, SAMP8 animals show elevated oxidative stress, chronic inflammation, altered microglia, decreased neurogenesis, and synaptic and neuronal dysfunction, among other symptoms (Tanisawa et al., 2013; Griñan-Ferré et al., 2016; Yanai and Endo, 2016; Díaz-Moreno et al., 2018). Furthermore, the SAMP8 model develops neurodegenerative changes with age, and has been proposed as a relevant model for the study of age-related diseases such as late-onset Alzheimer’s disease (Morley, 2002; Pallas et al., 2008; Akiguchi et al., 2017). Therefore, SAMP8 stands as a highly useful tool for studying the mechanisms of brain aging and exploring potential therapeutic interventions.

Astrocytes are the most abundant cell type in the brain (Miller, 2018). They fulfil fundamental functions in brain homeostasis, such as blood-brain barrier (BBB) maintenance, immune signalling, ionic level control, neurotrophin secretion, synapse formation and neurotransmitter recycling (Chen et al., 2006; Phatnani & Maniatis, 2015; Cheng et al., 2016; Miller, 2018). Astrocyte senescence has been proposed to contribute to the age-related loss of astrocyte function (Cohen & Torres, 2019; Gorgoulis et al., 2019). However, research aimed at uncovering the impact of astrocyte senescence on the synaptic loss associated with aging and neurodegenerative diseases is very scarce. Here, we have taken advantage of the SAMP8 model to specifically interrogate the contribution of astrocyte senescence to synapse loss in the aged hippocampus, a paradigmatic region for synaptic plasticity studies in the context of learning and memory. Taking advantage of the SA-β-galactosidase staining, we show that senescence increases in astrocyte-enriched layers of the SAMP8 hippocampi and in acutely isolated SAMP8 hippocampal astrocytes. We also employ two methodologies for exploring the synaptogenic capacity of senescent astrocytes: i) isolation of adult hippocampal astrocytes from SAMP8 and SAMR1 based on the astroglial cell surface marker ACSA-2, followed by primary culture in defined medium, and ii) differentiation of hippocampal neural stem cells (NSCs) derived from adult SAMP8 and SAMR1 mice into astrocytes. We characterize the astrocytes using a variety of senescence markers and assess their functionality in synaptogenesis assays, employing primary hippocampal neuronal cultures. Our results show that hippocampal SAMP8 astrocytes progressively accumulate senescence marks, downregulate the production of the pro-synaptogenic signal thrombospondin-1 (TSP-1) and precociously lose their synaptogenic activity during aging. The decrease in TSP-1 levels parallels the excitatory synapse reduction observed in all astrocyte-enriched layers of the SAMP8 hippocampus.

## Experimental Procedures

### Animals

The accelerated senescence SAMP8/TaHsd model and the senescence-resistant control SAMR1/TaHsd strain (Takeda et al., 1981) were used in this study. Mice were purchased from Inotiv Inc. and bred in the animal facility of the Biomedicine Institute of Valencia. Experiments were performed in male young (2 months, 2-m), middle-aged (6 months, 6-m) and old (10 months, 10-m) animals. For hippocampal neuron culture, RjOrl:SWISS wild-type mice were purchased from Janvier Labs, and timed-pregnant mice (E18.5) were bred in the animal facility of the Biomedicine Institute of Valencia. Mice were sacrificed by cervical dislocation for primary cultures, and by anesthetic overdose prior to perfusion for immunohistochemistry techniques. Animals were bred under controlled temperature conditions, 12 h light/dark cycle and water and food *ad libitum*. All experimental procedures and handling were in accordance with the European Union Council guidelines (2010/63/EU) and the Spanish regulation (RD53/2013). Procedures were approved by the Ethics and Animal Welfare Committee (CEEA) of the Biomedicine Institute of Valencia and CSIC (CEEA references: 2023-VSC-PEA-0212 and 2024-VSC-PEA-0094).

### Primary culture of ACSA-2^+^ SAMR1 and SAMP8 astrocytes

2-m, 6-m and 10-m SAMR1 and SAMP8 mice were euthanized by cervical dislocation and hippocampi were dissected from the brain. Hippocampal cell suspension was obtained using the Adult Brain Dissociation Kit (Miltenyi, 130-107-677) in combination with the gentleMACS™ Octo Dissociator (Miltenyi, 130-095-937). ACSA-2^+^ cells (astrocytes) were isolated by magnetic cell separation (MACS) method following the manufacturer’s procedure and protocols (Miltenyi Biotec, 130-097-679). ACSA-2^+^ cells were then seeded (200.000 cells) in 24-well poly-D-lysine (Sigma-Aldrich, 27964-99-4) and laminin-coated plate (Sigma-Aldrich, L2020), as recommended by Miltenyi Biotec for primary astrocyte cultures from the adult brain. Astrocytes were maintained in the defined commercial medium AstroMACS (Miltenyi Biotec, 130-117-031) supplemented with 2 mM L-glutamine (Lonza, 17-605C) and 100 u/mL penicillin-streptomycin (Lonza, 17-603E). Astrocytes were maintained for 14 days *in vitro* (DIV) and every other day, half of the medium was replaced. On the final day, cells were fixed with 2% paraformaldehyde (PFA) (PanReac, 141451). The cultures were maintained in an incubator at 37 °C in controlled humidity at 5% CO_2_.

### Primary culture of hippocampal neurons from wild-type mice

For the primary culture of hippocampal neurons, 18.5-day old embryos (E 18.5) from wild-type mice were used. Embryos were decapitated and hippocampal cell suspension was obtained using the Neural Tissue Dissociation Kit (Miltenyi, 130-092-628) in combination with the gentleMACS™ Octo Dissociator (Miltenyi, 130-095-937). Neurons were isolated by MACS using the Neuron Isolation Kit (Miltenyi Biotec, 130-115-390). Neurons were seeded (150.000 cells) onto poly-D-lysine and laminin-coated 24-well plate and maintained for 11 DIV. Neurobasal medium (Thermo scientific, 21103049) supplemented with B-27 (1X) (Thermo scientific, 17504044), 2 mM L-glutamine and 100 u/mL penicillin-streptomycin was used. On the first day post-plating, AraC (2 µM) was added to reduce non-neuronal contamination. The cultures were maintained in an incubator at 37 °C in controlled humidity at 5% CO_2_.

At 11 DIV, neuron medium was totally replaced by ACM from Ast-Diff or half replaced by ACSA-2^+^ ACM during 3h. Gabapentin (GBP, GenoChem World, HY-A0057) and TSP-1 (Dismed, HY-P701325) dosages were based in Cheng et al., (2016) procedures. GBP (32 μM) was added to neurons 30 minutes before ACM treatment and maintained 3.5 hours with the neurons. TSP-1 (250ng/mL) was added directly diluted with the ACM during 3 hours. Finally, neurons were fixed with 2% paraformaldehyde (PFA) and synapses were quantified.

### Culture of differentiated astrocytes (Diff-Astrocytes) derived from SAMR1 and SAMP8 neural stem cells (NSCs)

SAMR1 and SAMP8 NSCs were isolated from the hippocampal dentate gyrus of 2-m mice and expanded in the presence of mitogens following Babu’s protocol (Babu et al., 2011). These cells were maintained in adherent culture with Neurobasal medium supplemented with B-27 (1X), L-glutamine (2 mM), penicillin-streptomycin (100 u/mL), fibroblast growth factor 2 (FGF2 20 ng/mL) (PeproTech, 100-18B) and epithelial growth factor (EGF 20 ng/mL) (PeproTech, AF-315-09). For cellular assays, experiments were conducted using comparable passage (P) number for both models, and cells were discarded at approximately P26. To obtain SAMR1 and SAMP8 Diff-Astrocytes, NSCs were seeded (150.000 cells) in differentiation medium, consisting of Neurobasal medium supplemented with B-27 (1X), L-glutamine (2 mM), penicillin-streptomycin (100 u/mL) and Fetal Bovine Serum (FBS at 5%) (Thermo scientific, 10100147) for a period of 8 to 11 days. On day 4, half of the medium was replaced with fresh differentiation medium. Cultures were maintained in an incubator at 37 °C in controlled humidity at 5% CO_2_. For immunofluorescence staining, 8 DIV cells were fixed on coverslips in 2% PFA for 10 minutes.

SAMP8 Diff-Astrocytes were co-transfected at day 11 with pcDNA3 mTSP1 and pMaxGFP, or with empty pcDNA3 and pMaxGFP as a control, using Lipofectamine 2000 (Fisher scientific, 15338-030). pcDNA3 mTSP1 was a gift from Paul Bornstein (Addgene plasmid # 12405; http://n2t.net/addgene:12405; RRID: Addgene_12405). 24 hours after the transfection, the medium was changed to remove Lipofectamine. Cells were processed for RNA extraction or astrocyte conditioned medium collection 2 days later.

### Astrocyte conditioned medium (ACM) collection and treatment

Diff-Astrocyte cultures were seeded in 24-well plates and maintained 11 DIV. The culture medium was then completely removed and replaced with fresh differentiation medium for a 24-hours conditioning. After this time, the conditioned medium was collected and centrifuged at 300 g for 5 min to remove cellular traces. ACM from Diff-astrocytes and transfected Diff-astrocytes was immediately applied for 3 hours to the hippocampal neurons (via full media change), or stored at-20° C.

On the other hand, ACM collection from primary ACSA-2^+^ astrocyte cultures required a different procedure. Astrocytes were seeded in 24-well plates and half of the AstroMACS medium was collected and replaced every 48 h, for a period of 14 days. The medium was collected from each well, pooled, and centrifuged at 300 g for 5 min to remove cellular traces. For neuronal treatment, half of the neuronal culture medium was replaced during 3 hours with ACSA-2^+^ SAMR1 or SAMP8 ACM.

### Senescence-associated β-galactosidase (SA-β-gal) reaction

The SA-β-gal reaction was performed following the protocol established by Debacq-Chainiaux et al., (2009) with minor modifications. This reaction was performed in the same way for hippocampal tissues and for cell culture models. For cell cultures, the cells were fixed with 2% formaldehyde and 0.2% glutaraldehyde, followed by 3 washes with 0.1 M phosphate buffer (PB). The β-galactosidase staining solution was then prepared and the samples incubated with it for 5 hours at 37° C with relative humidity. Primary antibodies were then applied to label astrocytes and nuclei with DAPI (Sigma-Aldrich, D9542). Finally, the number of SA-β-gal positive cells for the astroglial markers GLAST and ATP1B2 was quantified as a percentage of the total cells photographed per field. For tissue, the sections were completely immersed in the staining solution for 5 hours and subsequently photographed. Quantification was interpreted as the SA-β-gal signal intensity normalized to the quantification area (µm^2^).

### Immunocytochemistry of astrocytes and neurons

To perform immunofluorescence techniques, cells were pre-fixed with 2% PFA for 10 minutes, followed by washing with 0.1M PB. Subsequently, the samples were incubated at room temperature for one hour with a blocking buffer composed of 0.1 M PB, 10% FBS and 0.5% Triton X-100. The primary antibody was then diluted in this blocking solution. This incubation was performed at 4 °C for 24 hours. The following day, cells were incubated with the secondary antibody, diluted in 0.1 M PB, for 2 hours. DAPI was used to label nuclei. Primary antibodies: mouse anti-GLAST (1:50; Miltenyi Biotec, 130-119-161), rabbit anti-ATP1B2 (1:200; Alomone Labs, ANP-012), guinea pig anti-MAP2 (1:1000; Synaptic Systems, 188004), rabbit anti-VGLUT1 (1:300; GeneTex, GTX133148), mouse anti-PSD95 (1:500; Thermo scientific, MA1045), rabbit anti-GFAP (1:500; Sigma-Aldrich, G3893), mouse anti-TSP-1 (1:50; Santa Cruz, sc-59887) and guinea pig anti-S100β (1:500; Synaptic Systems, 287004). Secondary antibodies: Alexa Fluor 488 anti-mouse (1:500; Invitrogen, A21202), Alexa Fluor 555 anti-mouse (1:500; Invitrogen, A31570), Alexa Fluor 488 anti-rabbit (1:500; Invitrogen, A21206), Alexa Fluor 647 anti-rabbit (1:500; Invitrogen, A31573), Alexa Fluor 488 anti-guinea pig (1:500; Jackson, 706-545-148) and Alexa Fluor 647 anti-guinea pig (1:500; Jackson, 706-605-148).

### Immunohistochemistry of mouse hippocampal tissue

The brains were processed with a Leica VT 1200 vibratome. Serial 40 µm sections comprising the entire hippocampus were collected. Before immunostaining, the tissue was washed three times for 5 minutes with 0.1 M PB to remove azide from the storage medium. The tissue was then incubated at room temperature for 1 hour with a blocking buffer composed of 0.1 M PB, 10% FBS, and 0.5% Triton X-100. The primary antibody was then diluted in the blocking buffer, and the sections were completely immersed in the mixture and kept shaking at 4°C for 24 or 48 hours. The next day, the slices were then incubated for 2 hours with the secondary antibody diluted in 0.1 M PB, protecting them from light. DAPI was used to label nuclei. Primary antibodies: rabbit anti-VGLUT1 (1:300; GeneTex, GTX133148), mouse anti-PSD95 (1:500; Thermo scientific, MA1045), rabbit anti-GFAP (1:500; Sigma-Aldrich, G3893), mouse anti-TSP-1 (1:50; Santa Cruz, sc-59887) and guinea pig anti-S100β (1:500; Synaptic Systems, 287004). Secondary antibodies: Alexa Fluor 555 anti-mouse (1:500; Invitrogen, A31570), Alexa Fluor 488 anti-rabbit (1:500; Invitrogen, A21206), Alexa Fluor 647 anti-rabbit (1:500; Invitrogen, A31573) and Alexa Fluor 488 anti-guinea pig (1:500; Jackson, 706-545-148).

### Confocal imaging and image analysis

Cells and tissue were observed with a Leica TCS SP8 confocal microscope and LSM 980 confocal microscope with ZEISS Airyscan detector. For the SA-β-gal reaction in cells, the images were obtained on a Leica DM6 B Automatic Upright Microscope (Thunder) and for tissue a Leica Aperio Versa Digital Scanner was used. Tissue images were captured at 1024×1024 resolution with a 40x focal lens, tracing vertical maps to obtain representative areas of all hippocampal region. Cell photographs were taken with a 40x or 63x focal lens at 512×512 resolution, with at least 5 representative photographs of each coverslip. The colocalization synapse analysis was performed by neuronal immunostaining of pre-and post-synaptic markers (VGLUT1 and PSD95 respectively) and normalized to the total number of neurons (MAP2^+^) of the image, using the Puncta Analyzer plugin and protocol of analysis (Ippolito & Eroglu, 2010), with Fiji ImageJ software.

### Quantitative RT-qPCR

RNA was extracted from cell cultures using a column extraction kit (Cytiva, 25050071) and subsequently quantified using a NanoDrop spectrophotometer. cDNA was obtained using the PrimeScript RT Reagent Kit (Takara, RR037A), following manufacturer’s instructions. The qPCR reaction was performed using the TB Green Premix Ex Taq Kit (Takara, RR82LR) and a QuantStudio 5 thermal cycler (Applied Biosystems).

The specific forward and reverse oligonucleotides were as follows (5‘-3’): *Lmnb1*: (F) CAACTGACCTCATCTGGAAGAAC, (R) TGAAGACTGTGCTTCTCTGAGC; *Il1β*: (F) CAGGCAGGCAGTATCACTCA, (R) TAATGGGAACGTCACACACC; *Il6*: (F) CAAAGCCAGAGTCCTTCAGAG, (R) TGGTCCTTAGCCACTCCTTC; *Cdkn1a:* (F) GGCAGACCAGCCTGACAGAT, (R) TTCAGGGTTTTCTCTTGCAGAAG; *Cacna2d1* (F) CCAAATCTTCAGCCAAAGGAGC, (R) ATTGACAGGCGTCCATGTGT. mRNA expression levels were calculated using the 2-ΔΔCT method, according to Livak and Schmittgen (2001). *Sdha* was used as a housekeeping gene for normalization, (F) AGAGGACAACTGGAGATGGCATT, (R) AACTTGAGGCTCTGTCCACCAA.

### Glutamate uptake assay

Free glutamate in the culture medium from NSCs and SAMR1 or SAMP8 Diff-Astrocytes was quantified using the commercial Glutamate Assay kit, following the manufacturer’s instructions (Sigma-Aldrich, MAK004). Glutamate levels were normalized with a MTT (3-(4, 5-dimethylthiazolyl-2)-2, 5-diphenyltetrazolium bromide) assay for cellular viability.

### Measurements of TSP-1 expression by ELISA and Western blot

TSP-1 levels were determined in 10-m SAMR1 and SAMP8 hippocampi. Tissue lysates were obtained by diluting hippocampi with cold phosphate-buffered saline (PBS) supplemented with protease inhibitors (Roche) and subjecting the samples to freeze-thaw cycles and to mechanic disruption with a homogenizer. The lysates were centrifuged at 5000 g for 10 min at 4°C, and the supernatants were collected and used. TSP-1 protein measurements were determined using the Mouse Thrombospondin-1 ELISA kit (CliniSciences, orb1807782-48) following the manufacturer’s instructions. Sample protein content normalization was based on total protein concentration determined for each sample by Bradford assay.

SAMR1 and SAMP8 Diff-Astrocytes were washed with PBS and homogenized in lysis buffer, supplemented with a protease inhibitor cocktail (Roche). Cell lysates were centrifuged at 13200 rpm for 15 min at 4°C and supernatant protein concentration was determined in a Bradford assay (Pierce). 50μg of protein lysates were loaded and resolved on sodium dodecyl sulfate-polyacrylamide gel (SDS-PAGE), and transferred to a nitrocellulose membrane (Amersham). The membrane was stained with Ponceau for total protein determination, and blocked in 5% skim milk. Membrane was incubated overnight with anti-TSP1/2 (1:200, sc-133061, Santa Cruz Biotechnology) and anti-β-Actin (1:5000, A-5441, Sigma-Aldrich) primary antibodies. The anti-TSP1/2 primary antibody cross-reacts with TSP1 and TSP2, which are structurally and functionally related and form a distinct subfamily of thrombospondins. IRDye 680LT anti-mouse (1:5000, Licor, 925-68020) secondary antibody was used. Proteins were detected with LI-COR Odyssey and analyzed with Image Studio Lite.

## Statistical analysis

At least three different male SAMR1 and SAMP8 mice were used in all experiments. For *in vitro* culture assays, at least three independent experiments were performed. Error bars represent the standard error of the mean (SEM). All statistical analysis were performed using GraphPad Prism software version 10. Normality was assessed with Shapiro-Wilk test. Two-way ANOVA, One-way ANOVA, One-sample t-test (for RT-qPCR) and Student’s t-tests were performed. In all analyses, a p < 0.05 was considered significant. Probabilities are represented as: * p < 0.05, ** p < 0.01, *** p < 0.001 and **** p < 0.0001. Statistical analyses are indicated in the figure legends.

## Results

### The SA-β-gal senescence mark is upregulated early in astrocyte-enriched layers of the SAMP8 hippocampus

Previously, SA-β-gal staining has been found to increase in the hippocampus of aged rodents, being reported in pyramidal neurons of the CA3 and CA1 hippocampal regions (Geng et al., 2010; Piechota et al., 2016). However, the age-related accumulation of senescent SA-β-gal^+^ astrocytes throughout the hippocampus has been poorly explored. To evaluate the impact of astrocyte senescence on hippocampal function, SA-β-gal signal intensity was measured in brain sections of 2-, 6-and 10-month-old (2-m, 6-m and 10-m) SAMR1 and SAMP8 animals. We evaluated both the entire hippocampus and a selection of hippocampal layers with high astrocyte content, according to the percentage of cells expressing the astrocyte marker GFAP (Glial Fibrillary Acidic Protein) (Fig. 1A-B). When comparing animals of the same age, SA-β-gal signal intensity in the entire hippocampus of the SAMP8 strain increased significantly compared to the SAMR1 control strain (Fig. 1C-D, p < 0.0001). SA-β-gal signal intensity also increased from 6-m to 10-m in both strains (Fig. 1D). At an early age (2-m), all the evaluated astrocyte-enriched hippocampal layers (Stratum Oriens-SO, Stratum Radiatum-SR, Stratum Lacunosum Moleculare-SLM, Molecular Layer-MO and Polymorphic Layer-PO) showed signs of senescence in SAMP8 *vs.* SAMR1. For most layers, at 6-m and 10-m SA-β-gal signal intensity remained higher in SAMP8. Two-way ANOVA analysis indicated that the effect of the strain was significant for the entire hippocampus and for all the analysed layers, with an interaction between strain and age in the SR stratum (p < 0.05). The combination of SA-β-gal staining with immunofluorescence detection of the mature astrocyte markers S100β (S100 calcium-binding protein beta) and GFAP confirmed the presence of SA-β-gal^+^ astrocytes (Fig. 1E and Supplementary Fig. 1). Together, these data allow to conclude that the SA-β-gal senescence mark is precociously increased in SAMP8 hippocampal layers with high astrocyte content compared to SAMR1 and is detected in GFAP^+^S100β^+^ astrocytes.

**Figure 1.**
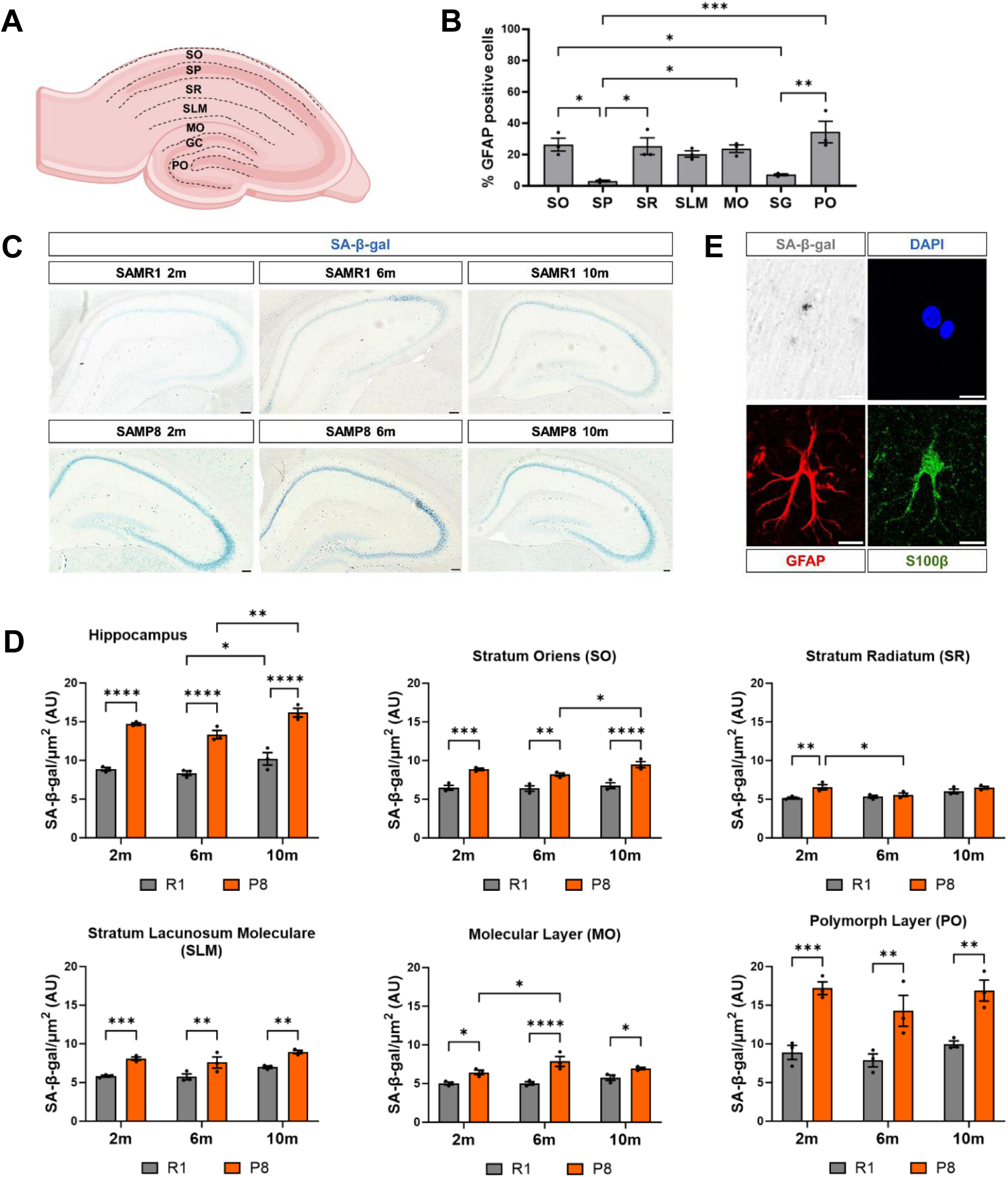
Senescence-associated β-galactosidase (SA-β-gal) activity in the hippocampus of SAMP8 mice show higher intensity at 2, 6 and 10 months than SAMR1 mice. (A) Diagram of the hippocampus cerebral layers: stratum oriens (SO), pyramidal layer (SP), stratum radiatum (SR), stratum lacunosum moleculare (SLM), molecular layer (MO), granule cell layer (GC) and polymorph layer (PO). (B) Percentage of astrocytes (GFAP^+^) in the different hippocampal layers. (C) Representative slices of hippocampus from SAMR1 and SAMP8 mice at 2, 6 and 10 months with the SA-β-gal staining. (D) Quantification of the SA-β-gal staining intensity in the whole hippocampus and in specific layers normalized to the area (µm^2^). (E) Representative SA-β-gal positive astrocyte from stratum radiatum layer with GFAP (red) and S100β (green) biomarkers. The slices have 40 µm of thickness. Three independent animals of each strain and age were analyzed (n=3). Data are presented as mean ± SEM. Two-way ANOVA test was performed. * p < 0.05, ** p < 0.01, *** p < 0.001 and **** p < 0.0001. Scale bar, C = 50 µm; E = 10 µm. Representation shown in A. was created with BioRender.com.

### Astrocytes derived from hippocampal SAMP8 neural stem cells enter senescence and fail to support synaptogenesis

Astrocytes are key regulators of neuronal synapse formation, maintenance and function, so we next designed experiments to explore the impact of hippocampal senescent astrocytes on hippocampal neuronal synaptogenesis. To model astrocyte senescence, we developed a simple and efficient *in vitro* method using hippocampal neural stem cells (NSCs) isolated from 2-m SAMR1 and SAMP8 animals (Fig. 2A). We employed Neurobasal medium supplemented with serum to stimulate astrocyte differentiation (McCarthy and De Vellis, 1980). Expression of the astrocyte markers GLAST (Glutamate and aspartate transporter), ATP1B2 (ATPase Na+/K+ transporting subunit beta 2), S100β and GFAP was monitored in the cultures (Supplementary Fig. 2). Quantifications showed that nearly 100% of the differentiated SAMR1 and SAMP8 cells were GFAP^+^S100β^+^ and GLAST^+^ATP1B2^+^ at 8 days in vitro (DIV), in all independent assays performed (Supplementary Fig. 2). Consistently, expression of the intermediate filament Nestin, a NSC and progenitor cell marker, was reduced upon differentiation (Supplementary Fig. 2). We also verified the acquisition of relevant astrocyte functions such as glutamate uptake, following incubation of the cells with 100 μM glutamate in HBSS. For both strains, the presence of free glutamate in the media was reduced in differentiated astrocytes compared to NSCs, indicating an increased reuptake upon differentiation (Supplementary Fig. 2). Thus, we concluded that astrocytes differentiated *in vitro* from hippocampal NSCs (Diff-Astrocytes hereafter) acquire mature astrocytic properties.

**Figure 2.**
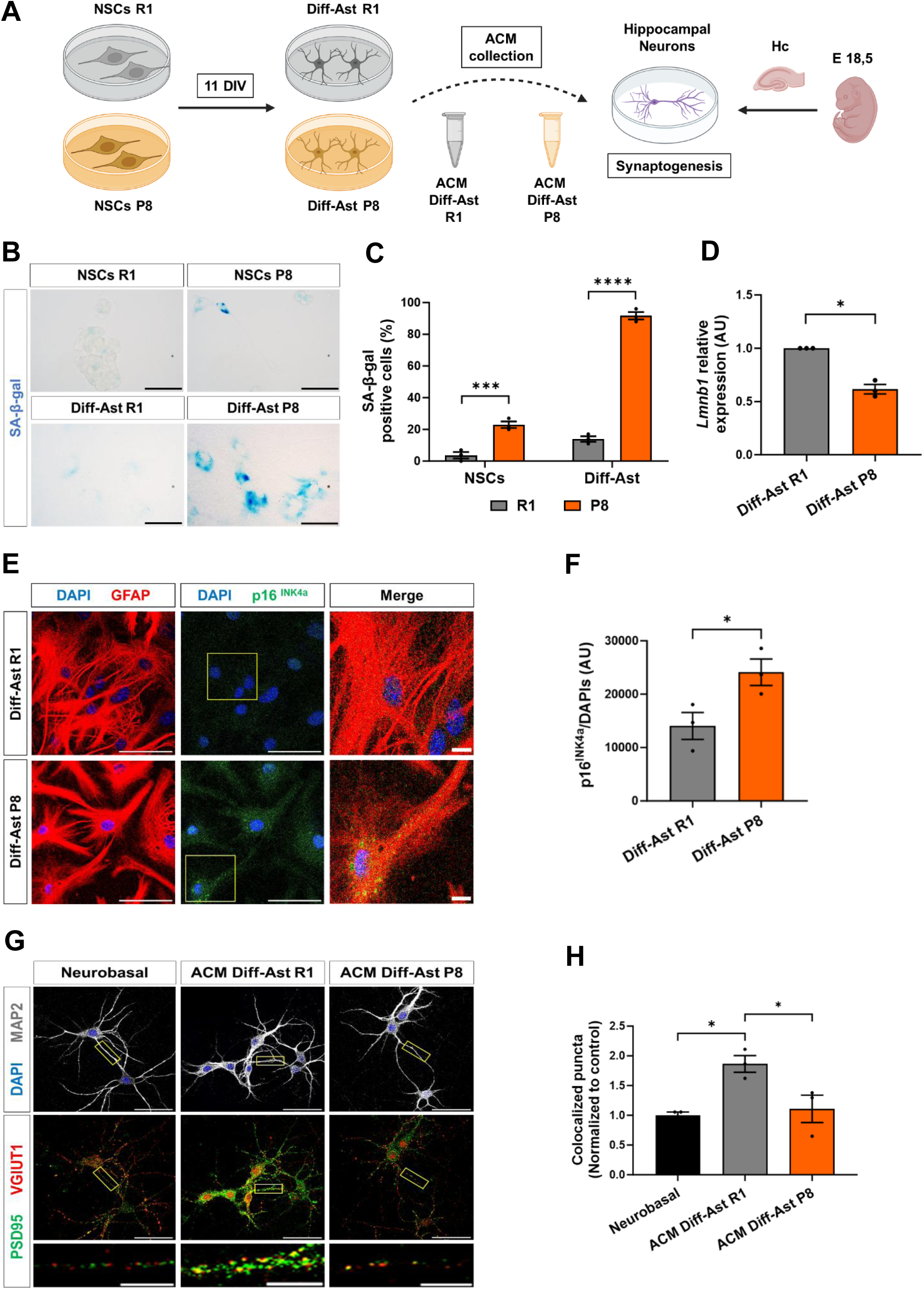
Differentiated astrocytes (Diff-Ast) derived from SAMP8 neural stem cells display hallmarks of senescence and synaptogenic dysfunction. (A) Schematic diagram of NSCs culture and differentiation to astrocytes. (B-C) SA-β-gal activity in NSCs and Diff-Ast SAMR1 (R1) and SAMP8 (P8) cultures. (D) RT-qPCR of *Lmnb1* (p = 0.013) in Diff-Ast SAMR1 and SAMP8 cultures. (E-F) Immunostaining of GFAP (red), p16INK4a (green), and p16INK4a quantification intensity in Diff-Ast SAMR1 and SAMP8 cultures. (G-H) Immunostaining of MAP2 (grey), PSD95 (green) and VGLUT1 (red), and quantification of excitatory pre-(VGLUT1) and postsynaptic (PSD95) vesicles colocalization in hippocampal neurons treated with ACMs from Diff-Ast SAMR1 and SAMP8 cultures. Three independent experiments were analyzed per mouse strain and cell type (n=3). Data are presented as mean ± SEM. One-way ANOVA Tukey’s multiple comparisons test was performed in (C) and (H). One-sample t-test was done in (D). Unpaired t-test in (F). * p < 0.05, ** p < 0.01, *** p < 0.001 and **** p < 0.0001. Scale bar: B, E and G = 50 µm; G = 10 µm (representative neurite). Representation shown in (A) was created with BioRender.com.

We next compared the SA-β-gal senescence mark in SAMR1 and SAMP8 NSCs and Diff-Astrocytes and quantified the percentage of positive cells. As expected, SAMP8 NSC and SAMP8 Diff-Astrocyte cultures showed a higher proportion of SA-β-gal^+^ cells compared to control SAMR1 cultures (p < 0.0001; Fig. 2 B-C). Interestingly, SAMP8 NSCs already displayed this senescence mark compared to SAMR1 NSCs, but the difference was greatly exacerbated upon NSC differentiation into astrocytes. Reduced Lamin B1 expression (p < 0.05; Fig. 2D) and increased p16^INK4a^ protein levels (p < 0.05; Fig. 2E-F), two other hallmarks of cellular senescence (Matias et al., 2021), were also detected in SAMP8 Diff-Astrocytes compared to SAMR1 Diff-Astrocytes by RT-qPCR and immunofluorescence, respectively. Expression of *Cdkn1a*/p21^CIP1^ and SASP genes further supported the acquisition of senescence (Supplementary Fig. 2).

Next, we evaluated if senescence influenced the synaptogenic capacity of the Diff-Astrocyte cultures. To this end, we established primary hippocampal neuronal cultures from wild-type embryos (E18.5) and tested the effect of astrocyte conditioned media (ACM) collected from SAMR1 and SAMP8 Diff-Astrocytes (Fig. 2A). Excitatory synapse formation was evaluated based on the immunofluorescence and co-localization image analysis of the pre-synaptic vesicle marker VGLUT1 (Vesicular Glutamate Transporter 1) and the post-synaptic marker PSD95 (Post-Synaptic Density protein 95), as previously described (Matias et al., 2021). ACM collected from SAMR1 Diff-Astrocytes promoted co-localization of the pre-and post-synaptic markers, demonstrating a positive impact on the generation of new synapses compared to control medium (p < 0.05; Fig. 2 G-H). Most importantly, ACM from SAMP8 Diff-Astrocyte cultures had no effect, demonstrating a loss of synaptogenic function in senescent astrocytes. Altogether, these results indicate that astrocytes differentiated *in vitro* from hippocampal SAMP8 NSCs enter senescence (as shown by the raise in SA-β-gal, p16^INK4a^ and p21^CIP1^, and the reduction in Lamin B1) and are deficient in promoting synaptogenesis compared to control (SAMR1) astrocytes, providing a suitable model to study the impact of astrocyte senescence in astrocyte-neuron communication.

### Acutely isolated ACSA-2^+^ hippocampal astrocytes from SAMP8 animals exhibit senescence marks and fail to support synaptogenesis

To further corroborate these findings, we next isolated astrocytes directly from the adult hippocampus of 6-m SAMR1 and SAMP8 mice employing magnetic cell separation (MACS), taking advantage of the astrocyte cell surface marker ACSA-2 (encoded by *Atp1b2*) (Fig. 3A). We analysed the acutely isolated ACSA-2^+^ cells by RT-qPCR for senescence and SASP markers. Lamin B1 expression was significantly decreased (p < 0.05; Fig. 3B), while IL-1β expression was upregulated (p < 0.05; Fig. 3B) in ACSA-2^+^ SAMP8 cells compared to ACSA-2^+^ SAMR1 cells. Hippocampal ACSA-2^+^ cells were then cultured *in vitro* in defined astrocytic growth medium (without serum) and were analysed by immunocytochemistry, with specific astrocyte markers and SA-β-gal staining (Fig. 3C). The vast majority of the hippocampal ACSA-2^+^ isolated cells expressed astrocyte markers upon culture and were double-positive for GLAST and ATP1B2 (average GLAST^+^ATP1B2^+^ cells ±S.E.M: 96.7±0.7% in SAMR1 cultures and 93.3±2.0% in SAMP8 cultures, n=3). On average, 9.3±4.5% of the GLAST^+^ATP1B2^+^ astrocytes were SA-β-gal^+^ in SAMR1 primary astrocyte cultures, and this percentage raised 3-fold in SAMP8 primary astrocyte cultures (33.2±4.1%, p < 0.05; Fig. 3C-D). Similar results were obtained when SAMR1 and SAMP8 astrocytes were isolated and cultured from 2-m and 10-m animals (Supplementary Fig. 1).

**Figure 3.**
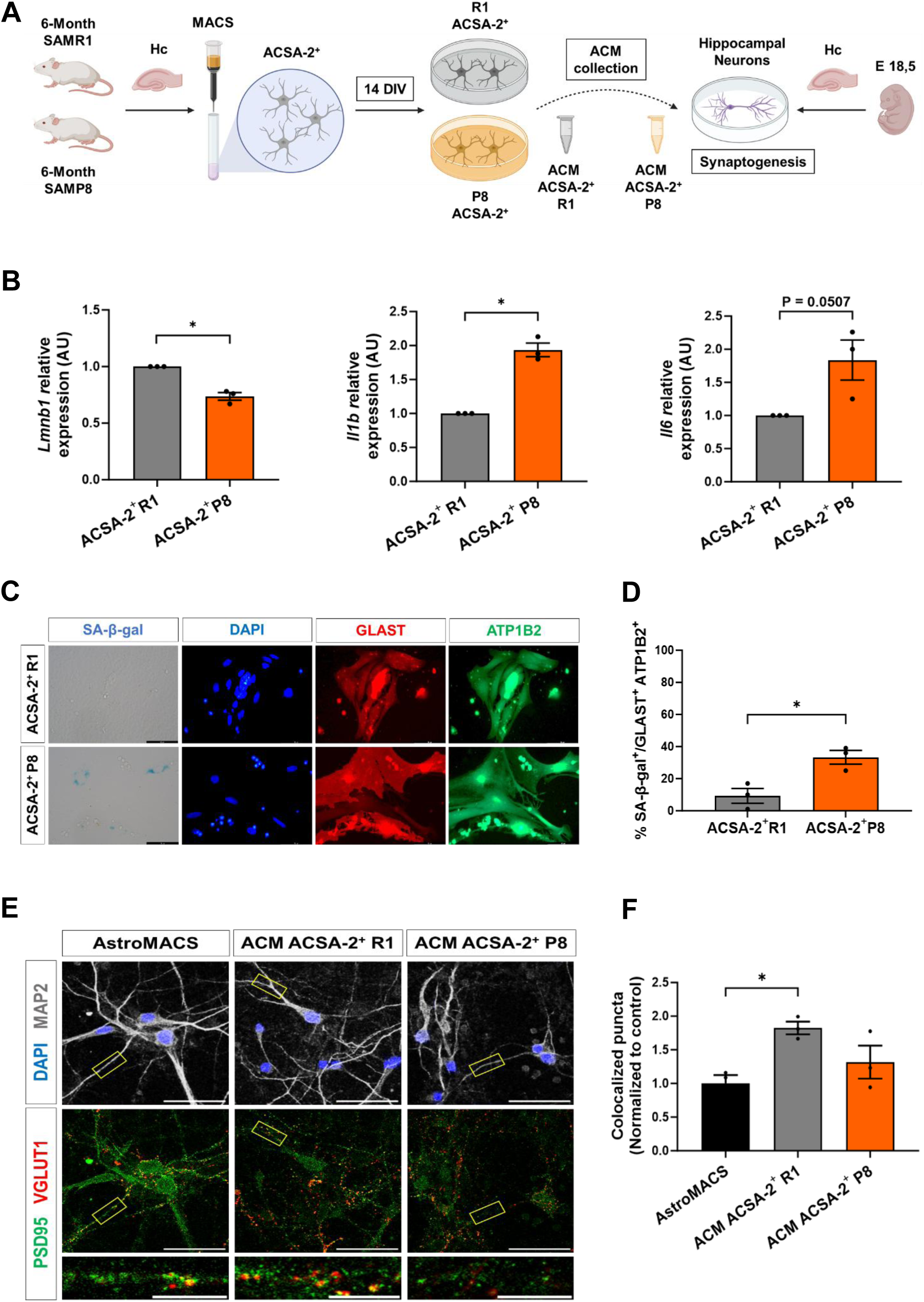
6-month-old ACSA-2 SAMP8 primary cultured astrocytes display signs of senescence and deficiencies in synaptogenic activity. (A) Experimental design of ACSA-2^+^ astrocytes isolation from the 6-month-old SAMR1 and SAMP8 hippocampus. (B) RT-qPCR of *Lmnb1* (p = 0.016), *Il1β* (p = 0.011) and *Il6* (p = 0.050) in ACSA-2^+^ SAMR1 and SAMP8 cells isolated from 6-m mice. (C-D) Representative images and quantification of SA-β-gal activity in ACSA-2^+^ primary cultures with astroglial GLAST (red) and ATP1B2 (green) markers after 14 days in culture. (E-F) Immunostaining of MAP2 (grey), PSD95 (green) and VGLUT1 (red), and quantification of excitatory pre-(VGLUT1) and post-synaptic (PSD95) vesicle co-localization in hippocampal neurons treated with ACMs from ACSA-2^+^ SAMR1 and SAMP8 astrocytes of 6-m. Three independent experiments were analyzed for each mouse strain (n=3). Data are presented as mean ± SEM. One-sample t-test was performed in (C). Unpaired t-test was done in (D). One-way ANOVA test was performed in (F). * p < 0.05. Scale bar: 50 and 10 µm. Representation shown in (A) was created with BioRender.com.

Next, we evaluated the synaptogenic activity of ACM collected from primary ACSA-2^+^ astrocytes, based on the co-localization of VGLUT1 and PSD95 in ACM-treated hippocampal neuronal cultures (Fig. 3E-F). SAMR1 ACSA-2^+^ ACM significantly promoted synapse formation compared to non-conditioned control medium (p < 0.05; Fig. 3F). In contrast, no synaptogenic effect was found when neurons were treated with SAMP8 ACSA-2^+^ ACM.

In summary, these results show that ACSA-2^+^ hippocampal astrocytes isolated from SAMP8 animals show senescence marks and are deficient in synaptogenic activity. While primary SAMR1 hippocampal astrocytes release factors into the ACM that promote the formation of new excitatory glutamatergic synapses in hippocampal neuronal cultures, this capacity is lost in primary SAMP8 hippocampal astrocytes, in line with the results obtained with senescent astrocytes differentiated from SAMP8 NSCs.

### Thrombospondin-1 expression is downregulated in senescent hippocampal SAMP8 ACSA-2^+^ astrocytes and in senescent astrocytes derived from SAMP8 neural stem cells

We then followed a candidate approach to uncover the identity of putative factors repressed in hippocampal senescent astrocytes that could account for the loss of synaptogenic function. Healthy brain astrocytes release molecules to the extracellular space that contribute to the formation of synapses (Ullian et al., 2004). Thrombospondin-1 (TSP-1, encoded by *Thbs1* gene) has emerged as an astrocyte-secreted protein with prominent effects on synaptogenesis and neuritogenesis during development (Cheng et al., 2016; Ikeda et al., 2010; Yu et al., 2008). We explored available bulk RNAseq databases *in silico* and confirmed the enriched expression of *Thbs1* in astrocytes among all other major cell types of the murine brain (Zhang et al. 2014, Figure 4A). Inspection of an RNAseq dataset from hippocampal astrocytes of postnatal, young, middle-age and old wild type animals highlighted the downregulation of *Thbs1* expression during aging (Clarke et al. 2018, Figure 4B). Given *Thbs1* gene expression is also reduced in astrocytes isolated from other brain areas during aging (Boisvert et al., 2018), we decided to delve deeper into *Thbs1*/TSP-1 expression in senescent astrocytes.

**Figure 4.**
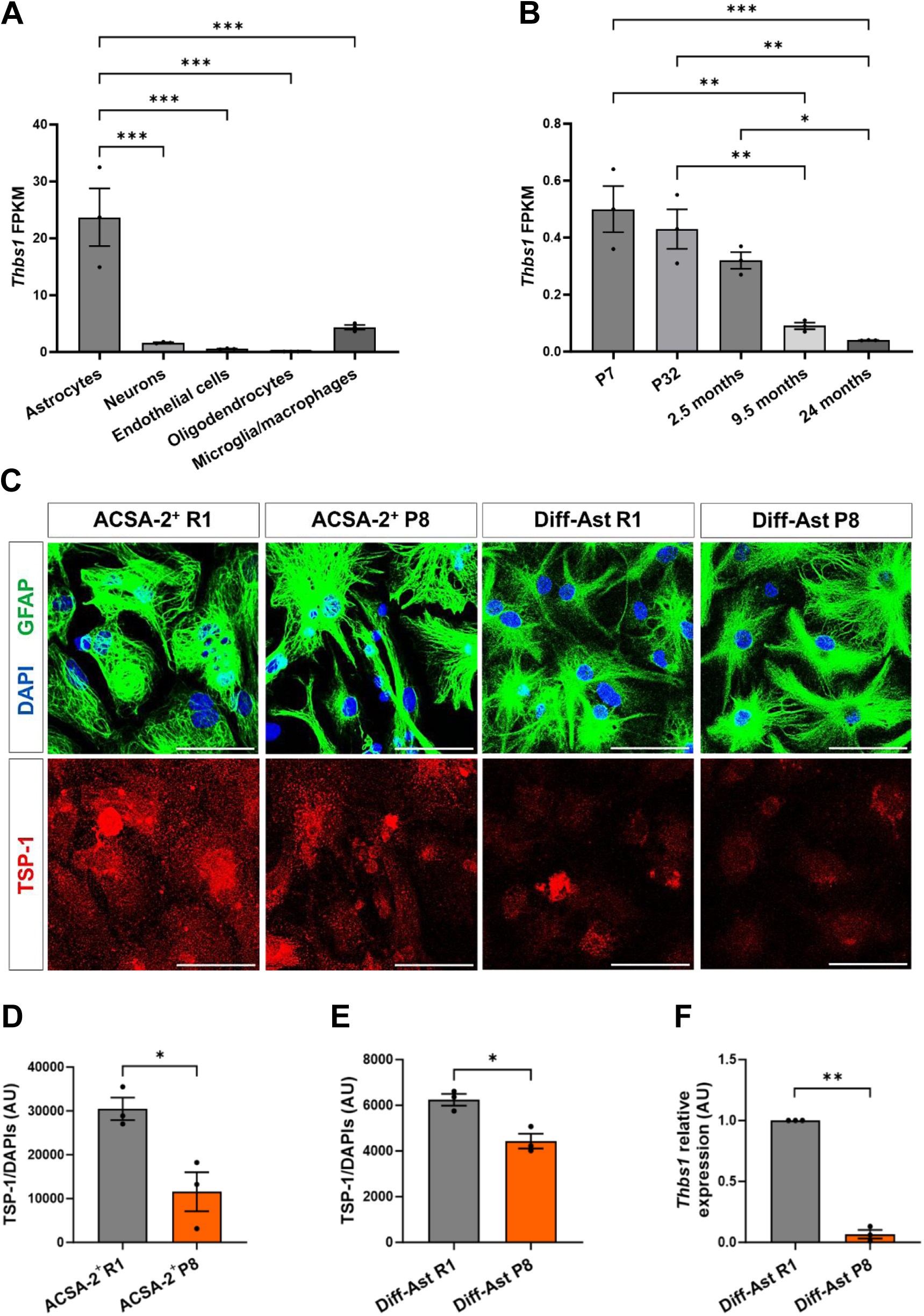
6-month-old ACSA-2^+^ SAMP8 primary cultured astrocytes and differentiated astrocytes derived from neural stem cells SAMP8 display loss in TSP-1 levels. (A) RNA-seq transcriptional profile of *Thbs1* gene in astrocytes, neurons, endothelial cells, oligodendrocytes and microglia in P7 (postnatal day) mice. (B) RNA-seq transcriptional profile of *Thbs1* gene at different stages of development and aging in astrocytes from mouse hippocampus. (C) Representative images of GFAP (green) and TSP-1 (red) immunostaining in ACSA-2^+^ primary astrocytes culture of the SAMP8 and SAMR1 strains (14 DIV) and in Diff-Ast SAMP8 and SAMR1 strains (11 DIV). (D-E) Quantification of TSP-1 protein show lower levels in SAMP8 ACSA-2^+^ astrocytes cultures (p = 0.021) and in Diff-Ast SAMP8 (p = 0.011) than in the control SAMR1 strain. (F) RT-qPCR of *Thbs1* (p = 0.001) demonstrates lower gene expression levels in Diff-Ast SAMP8 compared to the control line SAMR1. Three independent experiments per cell type were analyzed (n=3). Data are presented as mean ± SEM. Unpaired t-test was performed in (D-E). One-sample t-test was done in (F). * p < 0.05, ** p < 0.01, *** p < 0.001. Scale bar: 50 µm. Data in (A) and (B) were obtained from databases produced by the Barres lab (Zhang et al. 2014 and Clarke et al. 2018 available at https://brainrnaseq.org/).

We evaluated *Thbs1* mRNA and TSP-1 protein levels in senescent SAMP8 hippocampal astrocytes by RT-qPCR and immunocytochemistry. Our results demonstrate that primary SAMP8 ACSA-2^+^ astrocytes isolated from the hippocampus show significantly lower TSP-1 protein levels compared to SAMR1 ACSA-2^+^ astrocytes (p < 0.05; Fig. 4C-D). On the other hand, SAMP8 Diff-Astrocyte cultures also show decreased TSP-1 protein levels, as quantified by immunofluorescence (p < 0.05; Fig. 4C and 4E) and Western blot (Supplementary Fig. 3), and reduced *Thbs1* gene expression compared to SAMR1 Diff-Astrocytes (p < 0.01; Fig. 4F).

### Thrombospondin-1 expression and excitatory synapses decrease in the SAMP8 hippocampus

In order to further explore *Thbs1*/TSP-1 expression in the SAMP8 hippocampus *in vivo*, we measured *Thbs1* mRNA levels by RT-qPCR (Fig. 5A) and total TSP-1 protein levels by ELISA (Fig. 5B), in hippocampal tissue extracts from 10-m animals. Results confirmed the reduction in total *Thbs1* mRNA and TSP-1 protein content in SAMP8 hippocampus relative to SAMR1. This was further validated by immunofluorescence in hippocampal sections from SAMP8 relative to SAMR1 animals (Fig. 5C), where TSP-1 signal was found both co-localizing with the astrocytic marker GFAP and in the extracellular matrix surrounding the astrocytic processes. Confocal image analysis showed decreased TSP-1 levels in most astrocyte-enriched layers of the 10-m SAMP8 hippocampus compared to SAMR1 (Fig. 5D). Finally, we also analysed VGLUT1 and PSD95 co-localization in 10-m SAMP8 and SAMR1 hippocampal sections. As shown in Supplementary Fig. 4, we detected a significant decrease in co-localized VGLUT1 and PSD95 excitatory synaptic puncta, in most analysed SAMP8 hippocampal layers, and a significant correlation between TSP-1 signal intensity and puncta co-localization (Supplementary Fig. 4). These data support the notion that loss of TSP-1 expression in hippocampal senescent astrocytes may contribute to the reduction in excitatory synapses in SAMP8 animals.

**Figure 5.**
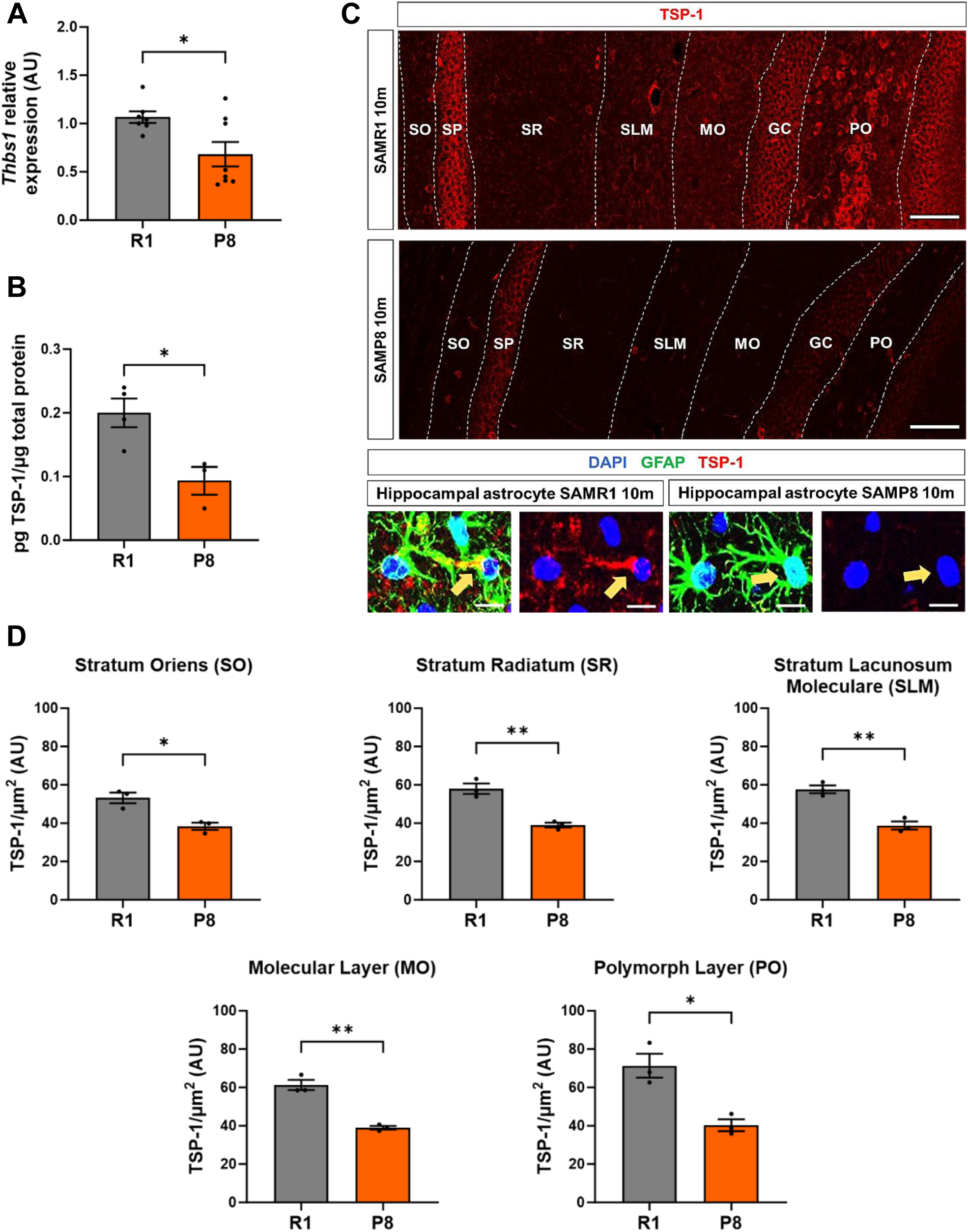
The hippocampus of 10-month-old SAMP8 mice shows deficiencies in TSP-1 levels. (A) RT-qPCR of *Thbs1* (p = 0.022) in SAMR1 (n = 7 independent animals) and SAMP8 (n = 8 independent animals). (B) TSP-1 quantification by ELISA test in 10-m SAMR1 (n = 4 independent animals) and SAMP8 (n = 3 independent animals) mice normalized to total protein levels. (C-D) Immunohistochemical analyses and TSP-1 intensity quantification in 10-m SAMR1 and SAMP8 hippocampus mice are shown. Representative astrocytes from the SLM layer in the hippocampus using GFAP (green) as an astroglial marker, and TSP-1 (red) are depicted magnified. TSP-1 signal intensity quantification was performed in each layer of the hippocampus and normalized to the quantification area (µm^2^). Three independent animals of each strain were analyzed for immunohistochemical analysis (n=3). Data are presented as mean ± SEM. Unpaired-t test was performed. * p < 0.05 and ** p < 0.01. Scale bar: 100 µm; 10 µm (representative astrocyte).

### TSP-1 released by healthy SAMR1 astrocytes is a major contributor to excitatory synaptogenesis through α2δ-1 receptor signalling

We next designed a functional assay to explore whether the synaptogenic effect induced by SAMR1 astrocytes is related to TSP-1 secretion. For this purpose, we employed Gabapentin (GBP), a pharmacological antagonist that blocks TSP-1 signalling through its neuronal receptor, the voltage-gated calcium channel (VGCC) subunit α2δ-1 (Cheng et al., 2016; Eroglu et al., 2009). The α2δ-1 receptor is encoded by the *Cacna2d1* gene, which is highly expressed in neurons throughout the CNS. The α2δ-1 receptor is enriched in hippocampal pyramidal neurons (Cole et al., 2005), and as shown by RT-qPCR, is expressed in the hippocampal neurons used in our cell culture assays (Supplementary Fig.5). As shown in Fig. 6, GBP completely blocked the pro-synaptogenic effect of SAMR1 Diff-Astrocyte ACM (p < 0.01; Fig. 6A and 6C). Similarly, GBP inhibited the positive effect of the SAMR1 ACSA-2^+^ ACM on the formation of new synapses (p < 0.05; Fig. 6B and 6D). No significant differences were found in GBP-treated cultures exposed to SAMP8 Diff-Astrocyte ACM or SAMP8 ACSA-2^+^ ACM.

**Figure 6.**
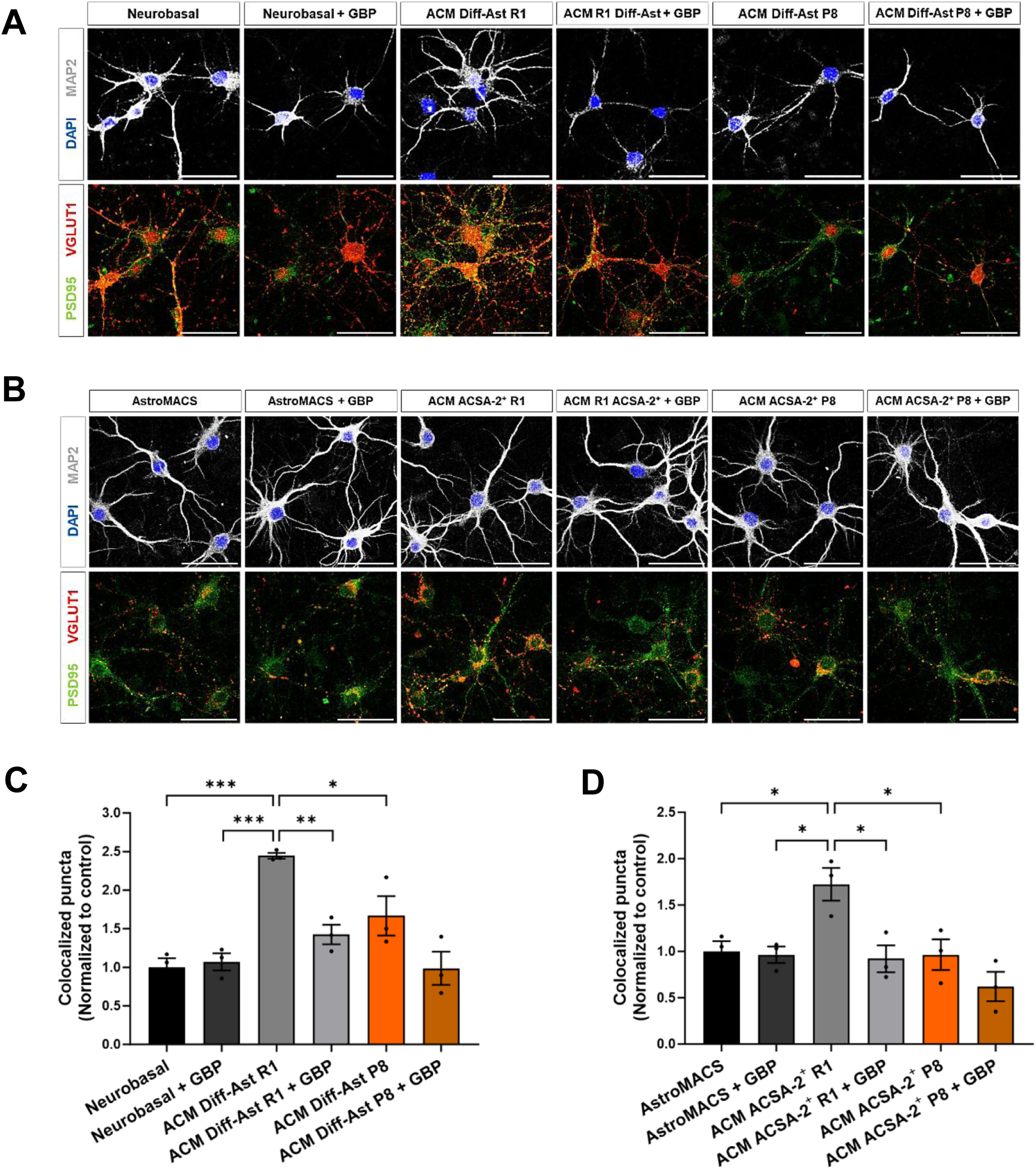
**The antagonistic competitor gabapentin (GBP) blocks the synaptogenic effect of SAMR1 ACMs**. (A and C) Immunostaining of MAP2 (grey), VGLUT1 (red) and PSD95 (green), and quantification of excitatory pre-(VGLUT1) and postsynaptic (PSD95) vesicles colocalization in hippocampal neurons treated with ACMs from Diff-Ast SAMR1 and SAMP8 with or without GBP 32 µM. (B and D) Immunostaining of MAP2 (grey), VGLUT1 (red) and PSD95 (green), and quantification of excitatory pre-(VGLUT1) and postsynaptic (PSD95) colocalization vesicles in hippocampal neurons treated with ACMs from ACSA-2^+^ SAMR1 and SAMP8 of 6-m mice with or without GBP 32 µM. Three independent experiments were analyzed per cell type and experimental condition. Data are presented as mean ± SEM. One-way ANOVA Tukey’s multiple comparisons test was performed. * p < 0.05, ** p < 0.01 and *** p < 0.001. Scale bar: 50 µm.

In summary, these results allow us to establish that the positive effect on the formation of new excitatory synapses triggered by healthy SAMR1 Diff-Astrocytes and SAMR1 ACSA-2^+^ primary hippocampal astrocytes is related to the presence of high TSP-1 levels in the ACM, which signals through the neuronal receptor α2δ-1.

### TSP-1 supplementation recovers the synaptogenic activity of senescent astrocyte conditioned medium

Finally, we explored whether *in vitro* treatment with purified recombinant TSP-1 restored the loss of synaptogenic activity of senescent SAMP8 astrocytes. Supplementing SAMP8 Diff-Astrocyte ACM with TSP-1 was sufficient to recover glutamatergic excitatory synapse formation in primary hippocampal neuronal cultures (p < 0.05; Fig. 7A-B). As a control, we further demonstrated that the pharmacological antagonist GBP blocked the TSP-1 effect on the co-localization of VGLUT1 and PSD95 pre-and post-synaptic markers (Supplementary Fig.5). Overexpression of the mouse *Thbs1* gene in SAMP8 differentiated astrocytes (Supplementary Fig. 5), followed by treatment of neuronal cultures with the resulting conditioned medium (Fig. 7C-D) showed that restoring TSP-1 expression suffices to rescue synaptogenic activity in senescent SAMP8 astrocytes.

**Figure 7.**
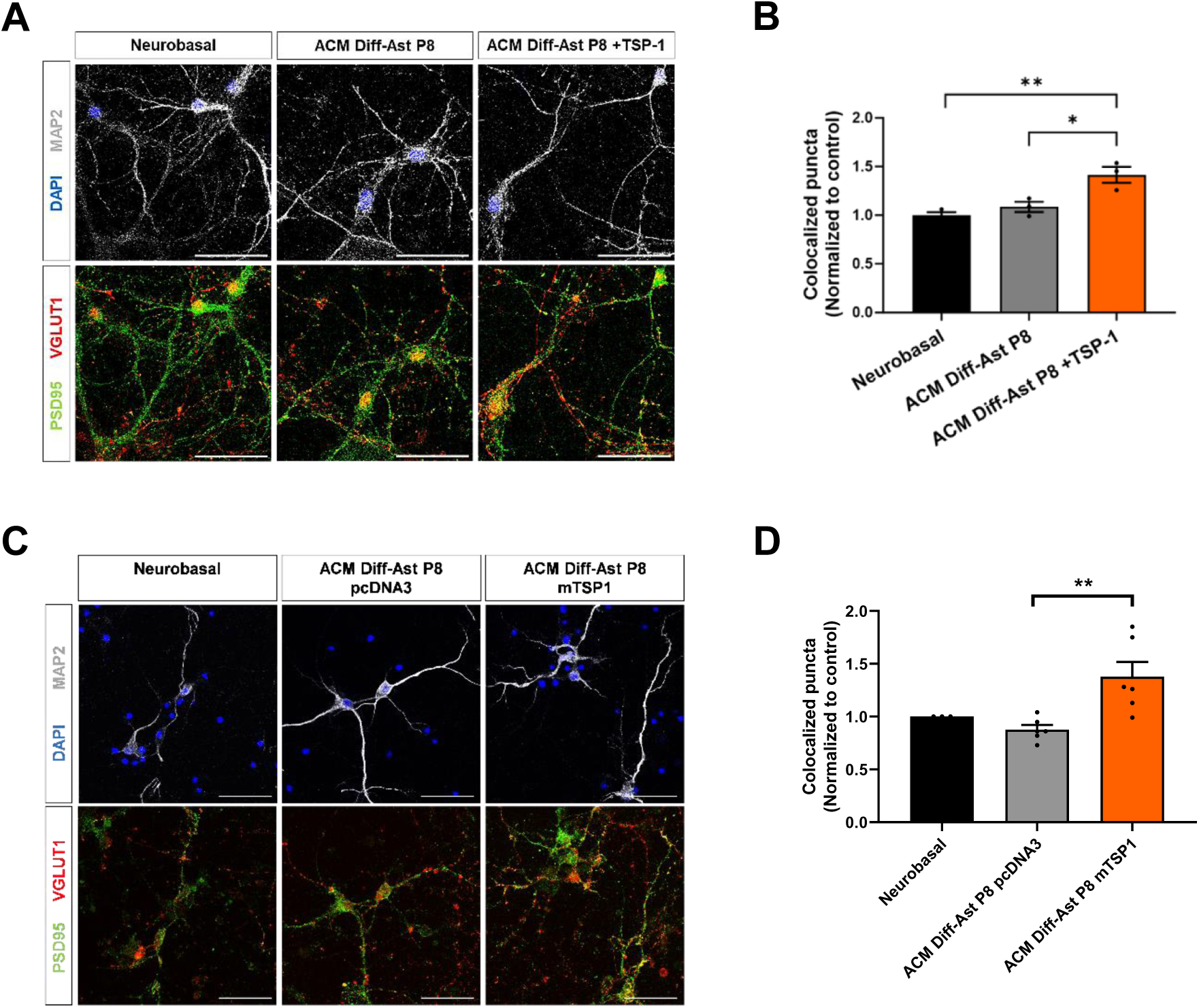
TSP-1 rescues the synaptogenic function of Diff-Ast SAMP8 ACM. (A-B) Immunostaining of MAP2 (grey), VGLUT1 (red) and PSD95 (green), and quantification of excitatory pre-(VGLUT1) and postsynaptic (PSD95) vesicles colocalization in hippocampal neurons treated with ACM from Diff-Ast SAMP8 with or without TSP-1 (250 ng/mL) supplementation. (C-D) Immunostaining of MAP2 (grey), VGLUT1 (red) and PSD95 (green), and quantification of excitatory pre-(VGLUT1) and postsynaptic (PSD95) vesicles colocalization in hippocampal neurons treated with ACM from transfected Diff-Ast SAMP8 overexpressing *mThbs1* and GFP, or pcDNA3 and GFP as a control. Three independent experiments were analyzed per cell type and experimental condition. Data are presented as mean ± SEM. One-way ANOVA Tukey’s multiple comparisons test was performed. * p < 0.05 and ** p < 0.01. Scale bar: 50 µm.

## Discussion

In this study, we employed the accelerated aging mouse model SAMP8 to characterize the accumulation of senescent astrocytes in the aged hippocampus and study the functional impact of senescence on astrocyte-neuron communication and excitatory synapse formation. Astrocytes have a variety of biological functions, including the regulation of new synapse formation (Verkhratsky & Nedergaard, 2018). This function is mainly mediated by astrocyte-secreted protein factors (Baldwin & Eroglu, 2017). Astrocytes regulate the formation of different types of synapses, such as glutamatergic (Ullian et al., 2001), GABAergic (Elmariah et al., 2005; Hughes et al., 2010), glycinergic (Cuevas et al., 2005), and cholinergic synapses (Reddy et al., 2003; Cao & Ko, 2007). Here, we confirmed the accumulation of SA-β-gal^+^ hippocampal astrocytes in SAMP8 *in vivo*, both in histological sections and in isolated astrocytes expressing the surface antigen ACSA-2. The raise in senescent astrocytes was accompanied by a reduction at 10 months of age in the co-localization of the pre-and post-synaptic excitatory glutamatergic markers, VGLUT1 and PSD95, a proxy of synapse number. Using astrocyte cultures and ACMs, we focused on evaluating the impact of the senescent astrocyte secretome on synapse formation. We uncovered a reduction in TSP-1, an important astrocyte-secreted protein involved in the regulation of spine development and synaptogenesis, in senescent astrocytes. We showed that addition of exogenous TSP-1, or overexpression of *Thbs1*, is sufficient to rescue the synaptogenic deficiency of senescent astrocytes.

Previous studies showed that senescence can be induced *in vitro* in murine and human astrocytes, through long-term cell culture (Geng et al., 2010; Kawano et al*., 2012;* Matias et al., 2021; Saenz-Antoñanzas et al., 2024), oxidative stress treatment (Simmnacher et al., 2020) or X-irradiation (Limbad et al.,2020). Mimicking astrocyte aging *in vitro* through prolonged cell culture increases reactive oxygen species production and compromises the astrocyte’s neuroprotective capacity (Pertusa et al., 2007). A loss of synaptogenic function has been reported in cortical neonatal astrocytes from wild type mice that entered senescence after several weeks of *in vitro* cell culture, but the mechanism underlying this phenotype has remained elusive (Kawano et al., 2012; Matias et al., 2021). To our knowledge, our study is the first to demonstrate that synaptogenic defects characterize naturally senescent astrocytes directly purified from the aged brain, shedding light on the molecular basis of their lack of synaptogenic function and linking TSP-1 loss to astrocyte senescence.

For the isolation of senescent astrocytes from adult hippocampal tissue, we took advantage of the positive selection with antibodies against the astroglial surface antigen ACSA-2. Primary ACSA-2^+^ astrocyte cultures were established and maintained for 14 DIV in a defined serum-free medium. Despite the low yield of this type of culture, the approach is of great interest for studying the properties of astrocytes *ex vivo*, since in classical methodologies based on postnatal primary cultures expanded in the presence of serum, the characteristics of the astrocytes change and a reactive glial phenotype is induced (Caldwell et al., 2022). We found no significant differences in the purity of the ACSA-2 cultures between strains, with the astroglial markers GLAST and ATP1B2 reaching values close to 95% of positive cells in all cases. SA-β-gal staining demonstrated a higher proportion of senescent GLAST^+^ ATP1B2^+^ SA-β-gal^+^ astrocytes in SAMP8 cultures compared to SAMR1 cultures. We also confirmed other senescence markers in ACSA-2^+^ SAMP8 astrocytes by RT-qPCR (decreased expression of the gene encoding Lamin B1 and increased expression of the gene encoding IL-1β). Thus, this strategy allows to obtain naturally senescent astrocytes directly from the aged brain. Most importantly, it allowed us to prove that the glutamatergic synaptogenic function of primary senescent astrocytes is greatly impaired, as shown by the lack of effect of the ACSA-2^+^ SAMP8 ACM on wild type murine hippocampal primary neuronal cultures. Results were also confirmed when using ACM collected from *in vitro* differentiated SAMP8 astrocytes.

On the other hand, we derived SAMR1 and SAMP8 Diff-Astrocytes from hippocampal NSCs. This complementary strategy represents a simple and practical methodology for studying astrocyte senescence, allowing for large-scale preparations without the need to promote the acquisition of senescence through long-term culture or exposure to cell damaging agents. This approach is very efficient and reproducible, generating SAMP8 astrocyte monolayers highly enriched (∼90%) in SA-β-gal^+^ senescent astrocytes, with low Lamin B1 expression and elevated levels of p16^INK4a^. The raise in the cell cycle inhibitor p16^INK4a^ has been widely reported as a senescence hallmark (López-Otín et al., 2023). Transgenic mice have been developed to visualize and eliminate senescent cells from aged tissues based on this marker (p16-INK-ATTAC mice, Baker et al., 2011; p16-3MR mice, Demaria et al., 2014). Astrocytes derived from postnatal mice show higher levels of p16^INK4a^ after 30 to 35 DIV (Matias et al., 2021). In post-mortem brains of Parkinson’s disease patients, astrocytes accumulate senescence marks and p16^INK4a^ is increased in the substantia nigra (Chinta et al., 2018). Here we report that astrocytes differentiated from SAMP8 NSCs show higher p16^INK4a^ content compared to SAMR1 astrocytes after 11 DIV. Along the same lines, mRNA levels of other senescence markers corroborate the senescent state of SAMP8 Diff-Astrocytes. We detected a significant decrease in *Lmnb1* expression (encoding Lamin B1), a significant increase in *Cdkn1a* expression (encoding p21^CIP1^) and a trend towards the upregulation of the genes that code for IL-1β and IL-6, two secreted pro-inflammatory molecules related to SASP. Decreased Lamin B1 and increased IL-1β expression were confirmed in SAMP8 ACSA-2^+^ astrocytes. Interestingly, decreased Lamin B1 content has been proposed as a hallmark of astrocyte senescence in the hippocampal dentate gyrus of aged mice (Matias et al., 2021). Furthermore, a general reduction in Lamin B1 has been reported in the hippocampal granule cell layer of post-mortem human tissue from elderly individuals. In the same tissue samples, hippocampal astrocytes show reduced nuclear circularity, possibly related to the loss of Lamin B1 (Matias et al., 2021).

Studying the molecular basis of astrocyte-neuron communication requires analysing the protein composition of the astrocyte secretome. Leading groups in the field have demonstrated the poor coincidence between changes in astrocyte gene expression and secretome (Caldwell et al., 2022), so astrocyte transcriptomics are not informative enough if changes are not confirmed at the protein level. We focused on evaluating the synaptogenic effect of the protein factors secreted by healthy astrocytes and their alteration in senescent astrocytes. We observed that, for the formation of new excitatory glutamatergic synapses, SAMR1 control ACM is pro-synaptogenic, as expected based on the literature (Baldwin and Eroglu, 2017), while SAMP8 senescent ACM shows a loss of function. TSP-1 protein levels were significantly reduced in astrocytes from the SAMP8 strain compared to SAMR1. TSP-1 is a secreted matricellular protein from the thrombospondin family, described as a key factor in the regulation of structural synapse formation in several neuronal types (Christopherson et al., 2005; Eroglu et al., 2009; Xu et al., 2010; Cheng et al., 2016). TSP-1 and TSP-2 deficient mice have lower synaptic density in the cortex (Christopherson et al., 2005), so a similar phenotype could be expected in the hippocampus, taking into account that the α2δ-1 receptor is enriched in both cortical and hippocampal pyramidal neurons (Cole et al., 2005). Furthermore, α2δ-1 knockout animals show a significant reduction in the number, degree of maturation, and activity of excitatory synapses in the cortex (Risher et al., 2018), again highlighting the relevance of TSP-1/α2δ-1 signalling for synaptogenesis.

To support the role of hippocampal astrocyte-released TSP-1, we performed a functional assay using gabapentin (GBP), a drug used to treat epilepsy and neuropathic pain that acts as an antagonist competitor of the TSP-1 receptor α2δ-1 in neurons (Eroglu et al., 2009; Cheng et al., 2016). The synaptogenic activity of thrombospondin is mediated by its EGF-like domain, which binds to the extracellular domain of α2δ-1. The interaction is thought to cause a conformational change in α2δ-1 that in turn allows the recruitment of a synaptogenic signalling complex (Baldwin and Eroglu, 2017). At the post-synaptic level, this complex regulates synaptogenesis through the Rho GTPase Rac1, which in turn promotes the reorganization of the actin cytoskeleton (Risher et al., 2018). Our results clearly show that GBP blocks excitatory synapse formation in hippocampal neurons when added to the SAMR1 ACM.

TSP-1 promotes the formation of silent synapses (Christopherson et al. 2005), which contain NMDA receptors but lack functional AMPA receptors. These synapses are inactive under baseline conditions but can be rapidly “unsilenced” during activity-dependent processes such as long-term potentiation (LTP) (Malinow and Malenka, 2002; Kerchner and Nicoll, 2008). “Unsilencing” occurs via the insertion of AMPA receptors, thereby strengthening synaptic transmission and facilitating learning and memory. Therefore, silent synapses provide a reservoir of modifiable connections that act as a versatile substrate for experience-dependent synaptic plasticity.

Previous work shows that post-synaptically silent synapses persist in the hippocampus of both adult and aged rats (Sametsky et al., 2010). Although their prevalence declines slightly with age—potentially contributing to age-related cognitive impairment—they remain a viable target for therapeutic interventions aimed at enhancing cognitive function in aging (Kumar et al., 2007; Sametsky et al., 2010). Age-related cognitive decline has been attributed to dysregulated activity-dependent plasticity at functional synapses and/or impaired recruitment of AMPA receptors into silent synapses (Sametsky et al., 2010). Reduced TSP1 production in senescent animals may limit the available pool of silent synapses—at least in the accelerated aging SAMP8 mouse model—thereby possibly contributing to cognitive decline.

TSP-1 expression alterations have been previously related to synaptic defects in several pathologies, but the contribution of senescent astrocytes to these syndromes remains underexplored. Alzheimer’s disease (AD) is a devastating age-related neurodegenerative disorder that is associated with massive synapse loss in the early clinical phase (Masliah et al., 1991; Sze et al., 1997). TSP-1 expression is decreased in neurons of AD patients (Buee et al., 1992). Interestingly, thrombospondins accumulate in Ab plaques of post-mortem AD patient brain samples compared to control cases. Moreover, it has been suggested that increased thrombospondin expression in human brain evolution, may contribute to the cognitive abilities of the human species compared to chimpanzees and macaques, but also to the increased human vulnerability to AD and neurodegeneration (Cáceres et al., 2007). Loss of thrombospondin expression has been reported as well in astrocytes from Down’s syndrome, a disease associated with a reduction in synaptic density (Garcia et al., 2010). Abnormal hippocampal synaptogenesis also characterizes Fragile X Syndrome (FXS), the major cause of inherited mental retardation (Jawaid et al., 2018). Interestingly, astrocytes isolated from a mouse model of FXS display loss of TSP-1 protein expression. Alterations in excitatory synapse formation of FXS hippocampal neurons are prevented in the presence of healthy ACM or following the administration of TSP-1 (Cheng et al., 2016). Therefore, soluble TSP-1 has been proposed as a potential therapeutic target for mental retardation syndromes. Our results extend its potential use to treat age-related conditions that involve synaptic decline.

In summary, our results highlight the contribution of astrocyte senescence to neuronal microenvironment dysfunction during aging and, in particular, reveal a potential role in hippocampal synapse reduction. Our study also reinforces the use of the SAMP8 strain for the study of brain aging and provides two reproducible protocols for obtaining senescent astrocytes *in vitro*. Combining these methodologies, we demonstrate that TSP-1 released by healthy adult hippocampal astrocytes is a major contributor to synapse formation through α2δ-1 receptor signalling in neurons, and that reduced TSP-1 expression in senescent astrocytes impairs their synaptogenic capacity. Brain repair mechanisms aiming to recover synapse loss may take into account the poor performance of senescent astrocytes. Our findings also reinforce the use of senotherapeutics as promising strategies to mitigate the effects of aging on brain function.

## Acknowledgements

We thank all members of Helena Mira’s laboratory for fruitful discussions. We also thank technical assistance from Marife Cano and Rosa Viana, at the IBV Confocal Microscopy Core. This work was supported by a predoctoral fellowship from Ministry of Science, Technology, Knowledge and Innovation of Chile to S.E. and grants PID2022-141707NBI00 from Spanish Ministry of Science and Innovation and CIAICO/2022/74 from Generalitat Valenciana to H.M.

## Author contributions

S.E. and H.M. designed research; S.E., L.C.-C. and E.J. performed research and analyzed data; S.E. and H.M. wrote the first draft of the manuscript. All authors approved the manuscript.

## Abbreviations

CA1: Cornu Amonis 1
CNS: Central Nervous System
DIV: Days In Vitro
Diff-Astrocyte: Differentiated Astrocyte
FBS: Fetal Bovine Serum
GFAP: Glial Fibrillary Acidic Protein
GLAST: Glutamate-Aspartate Transporter
NSCs: Neural Stem Cells
SAMR1: Senescence Accelerated Mouse Resistant 1
SAMP8: Senescence Accelerated Mouse Prone 8
SASP: Senescence Associated Secretory Phenotype
SA-β-gal: Senescence-associated β-galactosidase

## Data Availability Statement

The data that support the findings of this study are available from the corresponding author upon reasonable request.

