## Supplementary material for "Astrocyte senescence impairs synaptogenesis due to Thrombospondin-1 loss": Figures S1-S5

### Figure S1

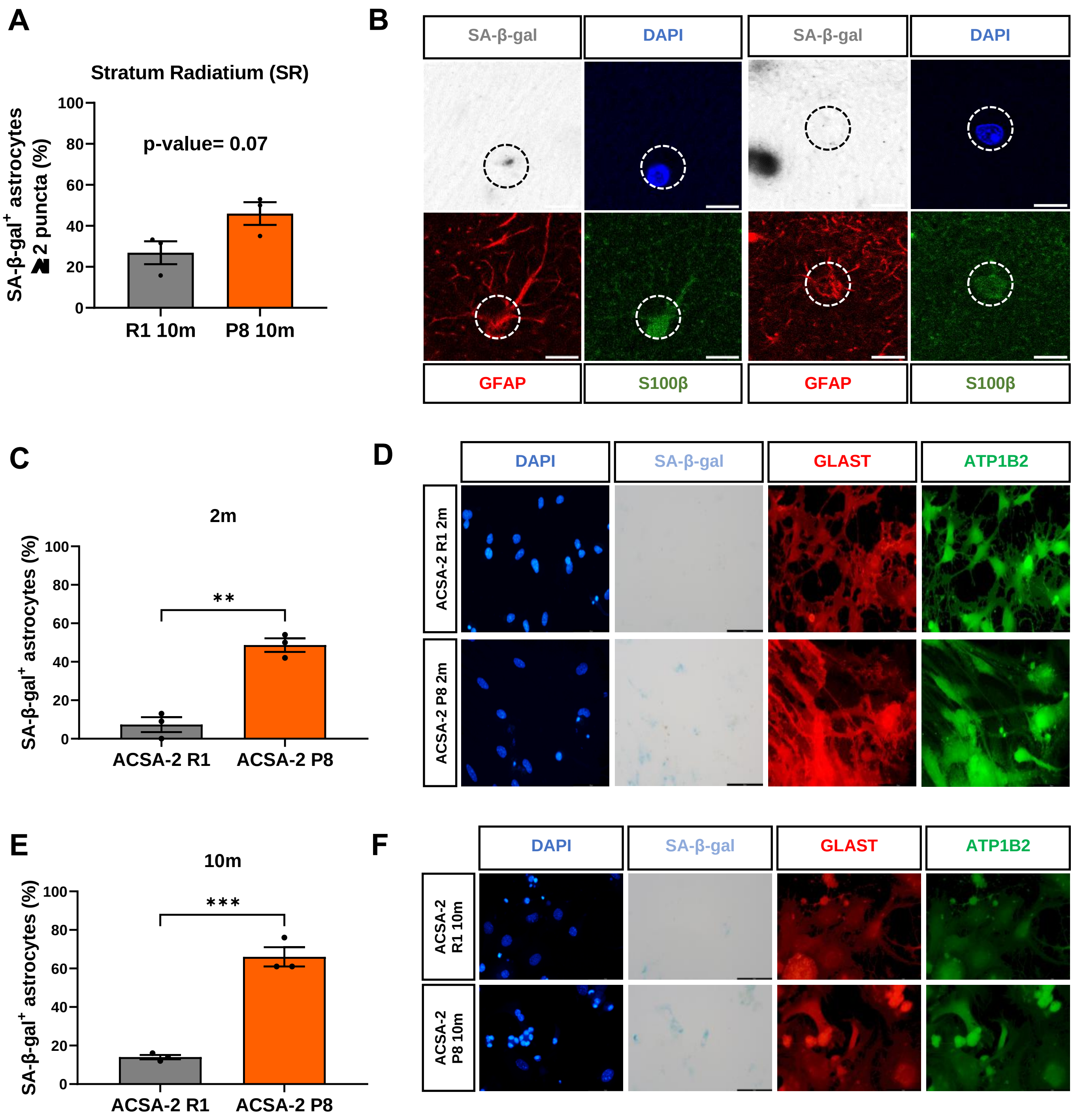
A
Stratum Radiatium (SR)
p-value= 0.07
SA-β-gal+ astrocytes ≥ 2 puncta (%)
R1 10m
P8 10m
B
SA-β-gal
DAPI
GFAP
S100β
C
2m
**
SA-β-gal+ astrocytes (%)
ACSA-2 R1
ACSA-2 P8
D
DAPI
SA-β-gal
GLAST
ATP1B2
ACSA-2 R1 2m
ACSA-2 P8 2m
E
10m
***
SA-β-gal+ astrocytes (%)
ACSA-2 R1
ACSA-2 P8
F
DAPI
SA-β-gal
GLAST
ATP1B2
ACSA-2 R1 10m
ACSA-2 P8 10m

**Supplementary Figure 1. SAMP8 hippocampi are enriched in SA-β-gal$^{+}$ astrocytes.** (A) Quantification of the percentage of astrocytes (GFAP$^{+}$/S100β$^{+}$) with two or more SA-β-gal puncta in the stratum radiatum of SAMR1 and SAMP8 mice at 10 months. (B) Representative SA-β-gal positive and negative astrocytes from stratum radiatum with GFAP (red) and S100β (green) biomarkers. The slices have 40 μm of thickness. (C) Percentage of SA-β-gal positive astrocytes (GLAST$^{+}$/ATP1B2$^{+}$) in ACSA-2 primary cultures of 2 months-old SAMR1 and SAMP8 mice. (D) Immunostaining of SA-β-gal (grey), GLAST (red) and ATP1B2 (green), in hippocampal astrocytes (ACSA-2$^{+}$) of 2 months-old SAMR1 and SAMP8 mice. (E) Percentage of SA-β-gal positive astrocytes (GLAST$^{+}$/ATP1B2$^{+}$) in ACSA-2 primary cultures of 10 months-old mice. (F) Immunostaining of SA-β-gal (grey), GLAST (red) and ATP1B2 (green), in hippocampal astrocytes (ACSA-2$^{+}$) of 10 months-old SAMR1 and SAMP8 mice. Three independent animals and primary cultures of each strain and age were analyzed (n=3). Data are presented as mean ± SEM. Unpaired t-test was performed. * $p < 0.05$, ** $p < 0.01$ and *** $p < 0.001$. Scale bar, C = 10 μm; E, G = 50 μm.

### Figure S2

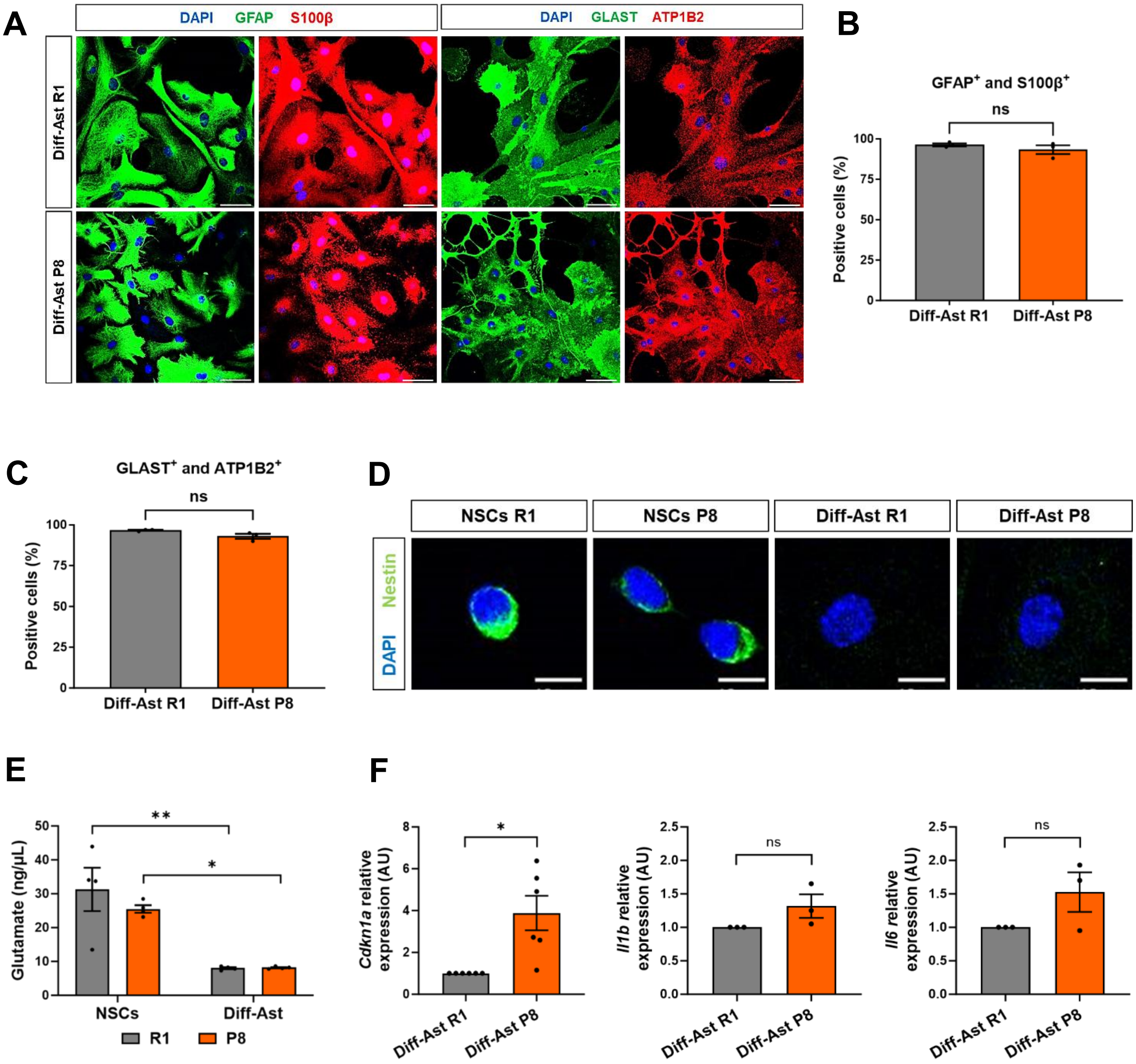

A
DAPI GFAP S100β
DAPI GLAST ATP1B2
Diff-Ast R1
Diff-Ast P8
B
GFAP+ and S100β+
ns
Positive cells (%)
100
75
50
25
0
Diff-Ast R1
Diff-Ast P8
C
GLAST+ and ATP1B2+
ns
Positive cells (%)
100
75
50
25
0
Diff-Ast R1
Diff-Ast P8
D
NSCs R1
NSCs P8
Diff-Ast R1
Diff-Ast P8
DAPI Nestin
E
Glutamate (ng/µL)
50
40
30
20
10
0
**
*
NSCs
Diff-Ast
R1
P8
F
Cdkn1a relative expression (AU)
8
6
4
2
0
*
Diff-Ast R1
Diff-Ast P8
Il1b relative expression (AU)
2.5
2.0
1.5
1.0
0.5
0.0
ns
Diff-Ast R1
Diff-Ast P8
Il6 relative expression (AU)
2.5
2.0
1.5
1.0
0.5
0.0
ns
Diff-Ast R1
Diff-Ast P8

**Supplementary Figure 2. Characterization and functionality of differentiated astrocytes (Diff-Ast) derived from neural stem cells.** (A) Representative images of SAMR1 and SAMP8 Diff-Ast using the astroglial markers GFAP (green) - S100β (red) and GLAST (green) - ATP1B2 (red). (B) Quantification of the percentage of $GFAP^{+}/S100\beta^{+}$ double positive cells in Diff-Ast. (C) Quantification of the percentage of $GLAST^{+}/ATP1B2^{+}$ double positive cells in Diff-Ast. (D) Representative images of SAMR1 and SAMP8 NSCs and Diff-Ast using the NSC marker Nestin (green). (E) Quantification of glutamate in proliferating NSCs at 2 DIV and Diff-Ast SAMR1 and SAMP8 at 8 DIV. Data represented free glutamate in the culture medium, normalized to cell viability. (F) RT-qPCR of *Cdkn1a* ($p = 0.017$), *Il1β* ($p = 0.214$) and *Il6* ($p = 0.217$) in Diff-Ast SAMR1 and SAMP8 at 11 DIV. At least three independent experiments per cell type were analyzed ($n \leq 3$). Data are presented as mean ± SEM. One-sample t-test was performed in (D). One-way ANOVA Tukey's multiple comparisons test was done in (E). * $p < 0.05$ and ** $p < 0.01$. Scale bar: 50 µm.

### Figure S3

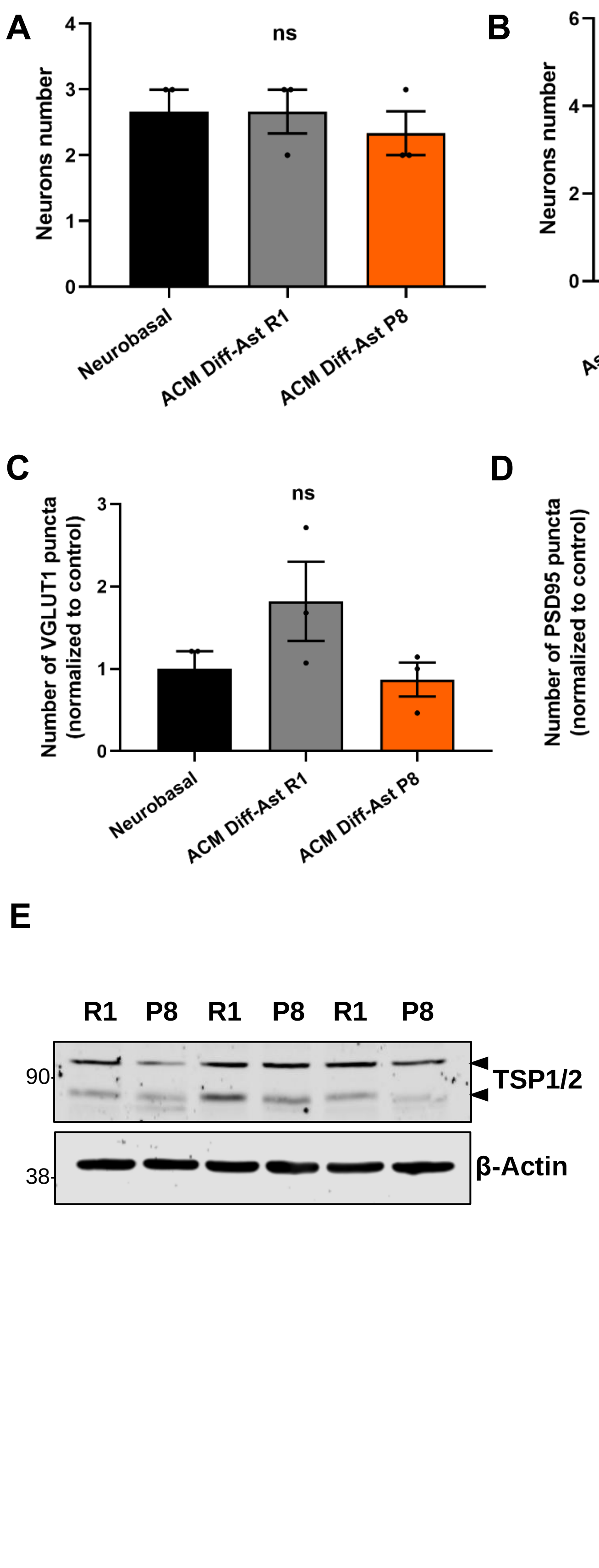
A
Neurons number
ns
Neurobasal
ACM Diff-Ast R1
ACM Diff-Ast P8
B
Neurons number
ns
AstroMACS
ACM ACSA-2+ R1
ACM ACSA-2+ P8
C
Number of VGLUT1 puncta (normalized to control)
ns
Neurobasal
ACM Diff-Ast R1
ACM Diff-Ast P8
D
Number of PSD95 puncta (normalized to control)
ns
Neurobasal
ACM Diff-Ast R1
ACM Diff-Ast P8
E
R1 P8 R1 P8 R1 P8
90
38
TSP1/2
β-Actin
F
TSP1/2 / Total protein (normalized to control)
*
Diff-Ast R1
Diff-Ast P8

**Supplementary Figure 3. Neuron survival, pre- and postsynaptic puncta are not affected upon ACM treatment, and TSP1/2 protein levels are decreased in SAMP8 differentiated astrocytes.** (A, B) $MAP2^+$ neuron counting in hippocampal cultures (8 random fields per culture) treated with ACM from SAMR1 and SAMP8 Diff-Astrocytes (A) and ACM from SAMR1 and SAMP8 $ACSA2^+$ primary astrocytes (B). (C, D) Quantification of pre- (VGLUT1) and postsynaptic (PSD95) vesicles in hippocampal neuron cultures treated with ACM from SAMR1 and SAMP8 Diff-Astrocytes. (E, F) Western blot showing TSP1/2 protein levels in three independent protein lysates from SAMR1 and SAMP8 Diff-Astrocytes, and their respective quantification. Three independent experiments were analyzed. Data are presented as mean ± SEM, and normalized to their respective controls in (C, D and F). One-way ANOVA Tukey's multiple comparisons test was performed in (A-D). Paired t-test was performed in (F). * $p < 0.05$.

### Figure S4

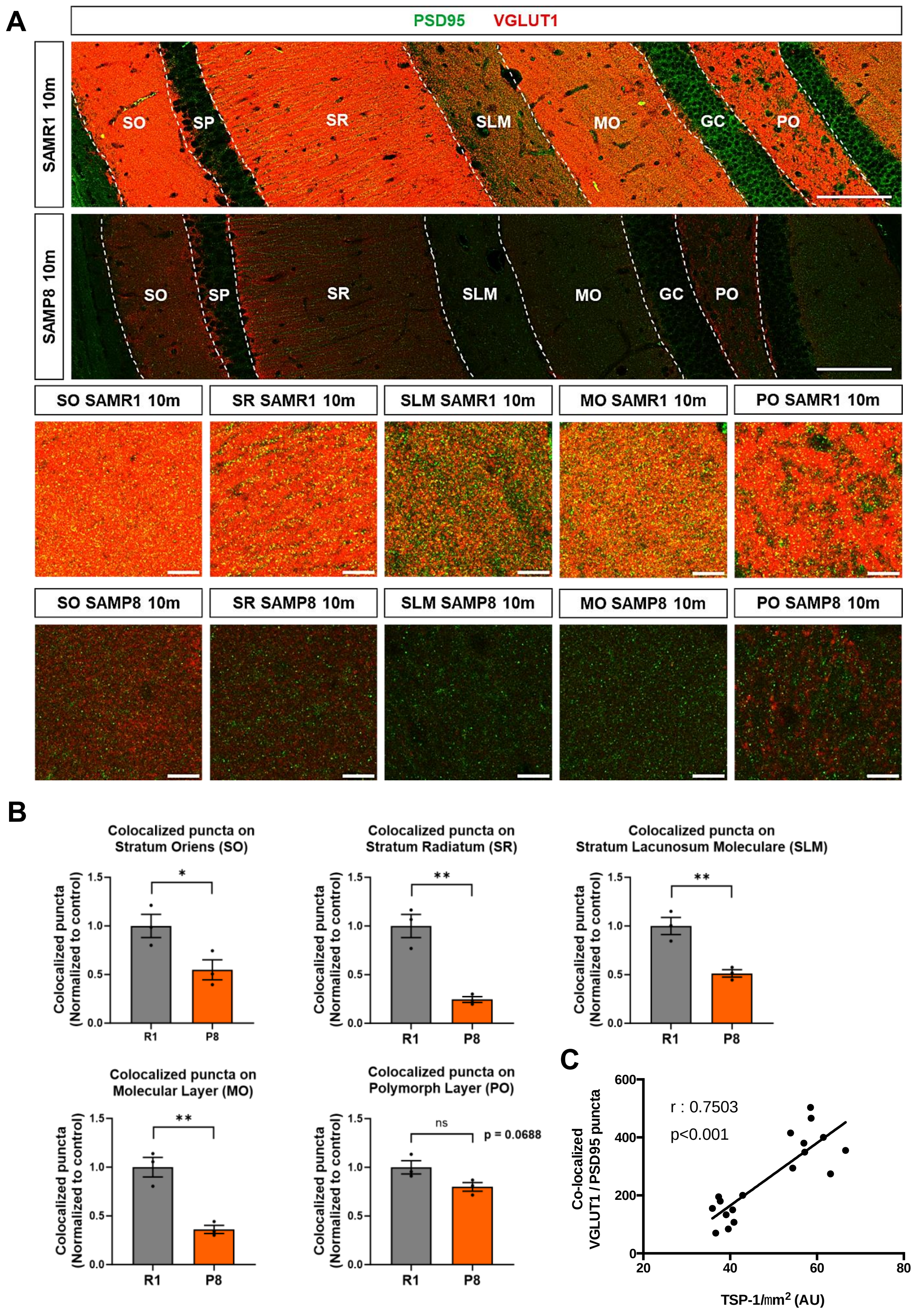
A
PSD95 VGLUT1
SAMR1 10m
SO
SP
SR
SLM
MO
GC
PO
SAMP8 10m
SO
SP
SR
SLM
MO
GC
PO
SO SAMR1 10m
SR SAMR1 10m
SLM SAMR1 10m
MO SAMR1 10m
PO SAMR1 10m
SO SAMP8 10m
SR SAMP8 10m
SLM SAMP8 10m
MO SAMP8 10m
PO SAMP8 10m
B
Colocalized puncta on Stratum Oriens (SO)
Colocalized puncta on Stratum Radiatum (SR)
Colocalized puncta on Stratum Lacunosum Moleculare (SLM)
Colocalized puncta on Molecular Layer (MO)
Colocalized puncta on Polymorph Layer (PO)
Colocalized puncta (Normalized to control)
R1
P8
ns
p = 0.0688
C
r : 0.7503
p<0.001
Co-localized VGLUT1 / PSD95 puncta
TSP-1/mm2 (AU)

**Supplementary Figure 4. Colocalization analysis shows a decrease in synapses in SAMP8 10-month-old hippocampal slices.** (A-B) Immunostaining and quantification of excitatory pre- (VGLUT1, red) and postsynaptic (PSD95, green) vesicles colocalization in hippocampal tissue of SAMR1 and SAMP8 at 10-m. Three independent animals of each strain were analyzed (n=3). (C) Correlation analysis of VGLUT1/PSD95 co-localized puncta, reflecting excitatory synapses, and TSP1 signal intensity normalized to the area ($\mu m^2$) in Stratum Radiatum, Stratum Lacunosum Moleculare and Molecular Layer. Three independent animals of each strain were analyzed (n=3). Spearman $r = 0.7503$ , Two-tailed $p < 0.0001$. Data are presented as mean ± SEM and normalized to SAMR1 mice. Unpaired t-test was performed. * $p < 0.05$, ** $p < 0.01$.

Figure S5

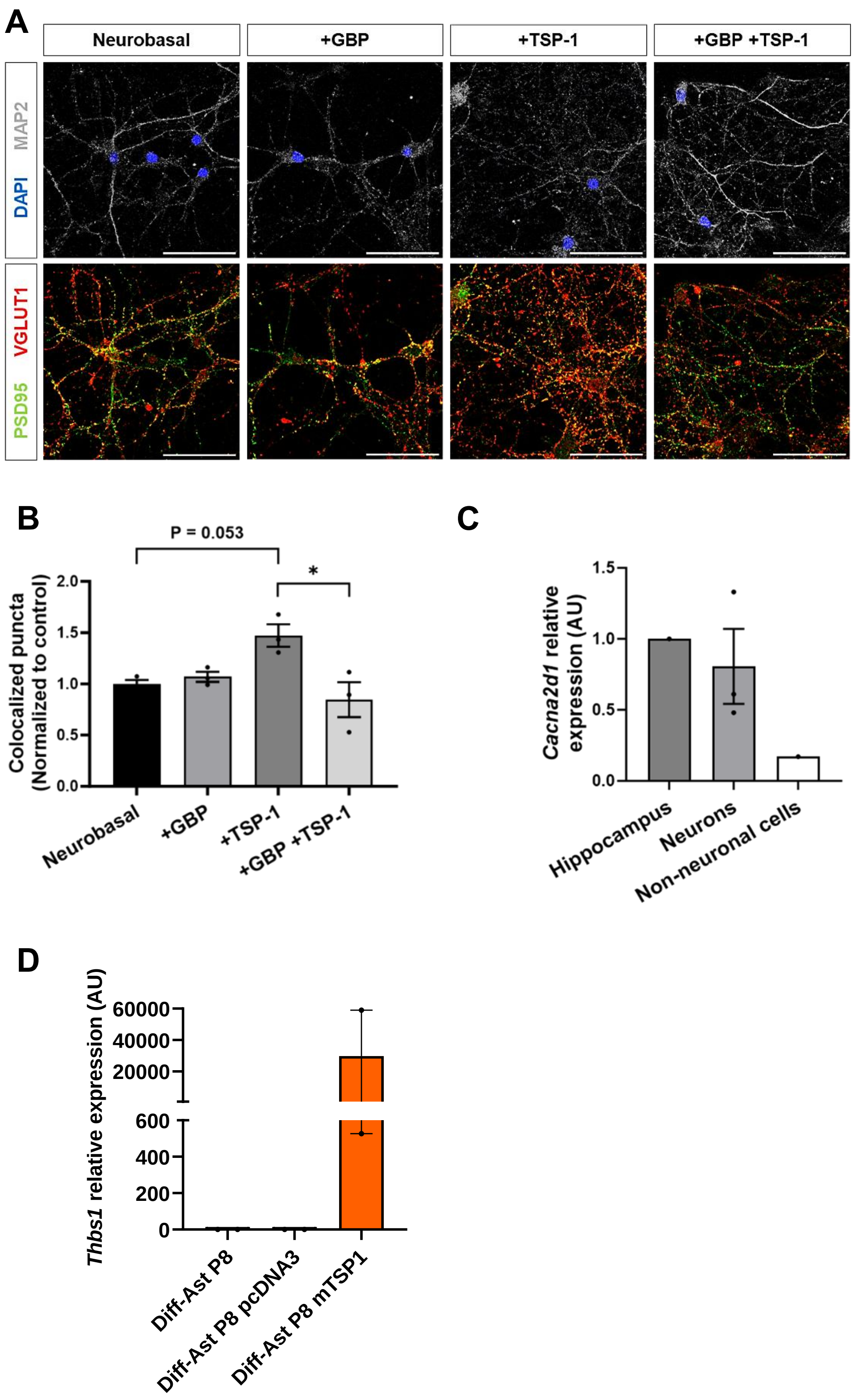
A
Neurobasal
+GBP
+TSP-1
+GBP +TSP-1
DAPI MAP2
PSD95 VGLUT1
B
P = 0.053
*
Colocalized puncta (Normalized to control)
2.0
1.5
1.0
0.5
0.0
Neurobasal
+GBP
+TSP-1
+GBP +TSP-1
C
Cacna2d1 relative expression (AU)
1.5
1.0
0.5
0.0
Hippocampus
Neurons
Non-neuronal cells
D
Thbs1 relative expression (AU)
60000
40000
20000
600
400
200
0
Diff-Ast P8
Diff-Ast P8 pcDNA3
Diff-Ast P8 mTSP1

**Supplementary Figure 5. The antagonistic competitor GBP shows a negative synaptogenic effect and TSP-1 rescues synaptic function in neurons.** (A-B) Immunostaining of excitatory pre- (VGlut1, red) and postsynaptic (PSD95, green) vesicles colocalization in hippocampal neurons of mice primary cultures. (C) RT-qPCR of *Cacna2d1* in neuronal and non-neuronal cells, normalized to whole hippocampus. (D) RT-qPCR of *Thbs1* in SAMP8 astrocytes, 3 days after transfection with pcDNA3.1 empty vector and pMaxGFP, or pcDNA3.1 mTSP1 and pMaxGFP (n=2). Three independent experiments per cell type were analyzed (n=3) in (B). Data are presented as mean ± SEM and normalized to the Neurobasal control medium in (B). One-way ANOVA Tukey's multiple comparisons test was performed in (B). * $p < 0.05$. Scale bar: 50 µm.
